# Implication of a rare variant in OPA1 in Cardiac Pathophysiology: From Cristae Remodelling to Contractile Dysfunction

**DOI:** 10.64898/2026.08.31.748193

**Authors:** Mohini Gupta, Amrita Mukhopadhyay, Ashok Kumar, Bhagyalaxmi Mohapatra

## Abstract

Optic Atrophy 1 (OPA1), an important inner mitochondrial membrane GTPase, regulates mitochondrial fusion, maintains cristae structure, calcium buffering, cellular bioenergetics, preserves mtDNA and controls apoptosis. Here we examined the role of OPA1 variants in DCM using whole-exome sequencing (WES) of 5 familial and 10 sporadic DCM cases. A rare *de novo* OPA1 variant, c.563C>T (p.Pro188Leu), was identified in a DCM patient, which is absent in 100 healthy controls as well as in the 1000 Genomes, IndiGenomes and GenomeAsia 100k databases while it showed very low MAF (0.000069) in GnomAD. Structural modelling predicted the variant to be highly deleterious and revealed marked conformational distortion of the mutant protein (RMSD = 3.5Å). Molecular docking further demonstrated enhanced accessibility of mutant OPA1 to mitochondrial protease OMA1, suggesting increased OPA1 proteolytic processing and a consequent increase in mitochondrial fragmentation.

Functional analysis in stable H9C2 cardiomyoblast cells, demonstrated significantly reduced OPA1 protein expression, extensive mitochondrial fragmentation in mutant-OPA1 expressing cells. The mutant protein caused significant reduction in mitochondrial membrane potential, ATP generation, and oxygen consumption rate (OCR), together with elevated cytosolic Ca² and reactive oxygen species (ROS) levels. qRT-PCR analysis further revealed depletion in mtDNA copy number and increased in expression of intrinsic apoptotic markers Caspase3, 9 and Bax/Bcl-2 ratio.

The above findings collectively highlighted the significant impact of the OPA1 mutation on mitochondrial dynamics and cellular health, suggesting a significant correlation with the pathogenesis of DCM. Collectively, these findings suggest that OPA1-mediated mitochondrial dysfunction represents a potential therapeutic avenue for the management of DCM.

## Introduction

Mitochondria serve as a central hubs of cellular metabolism, orchestrating not only ATP generation but also regulates many other cellular processes ssuch as apoptosis, calcium buffering, ROS production and intracellular signalling. Mitochondria are not static entities; rather, they are dynamic organelles that continuously undergo biogenesis, fusion–fission cycles, and selective degradation by mitophagy (Noone et al., 2022). Cardiomyocytes are exceptionally enriched in mitochondria, which constitute nearly one-third of their total volume, underscoring their crucial role in maintaining myocardial proliferation, contractile function, and metabolic homeostasis (Piquereau et al., 2013).

OPA1 (Optic Atrophy 1) gene was discovered to be associated with dominant form of optic atrophy (Delettre et al., 2000; Alexander et al., 2000). It is a dynamin-related GTPase residing in the inner mitochondrial membrane (IMM), where it is integral to mitochondrial fusion, as well as contributes to oxidative phosphorylation, Ca+2 homeostasis, mitophagy, and mitochondrial DNA stability (Olichon et al., 2003; Frezza et al., 2006; Barrera et al., 2016; MacVicar & Langer, 2016; Pernas & Scorrano, 2016). OPA1 facilitates IMM fusion by means of a GTP-coupled reaction and interaction with cardiolipin (DeVay et al., 2009; Ban et al., 2017). OPA1 activity is tightly regulated by alternative mRNA splicing and proteolytic processing, generating eight human isoforms with tissue-dependent abundance (Olichon et al., 2007). In the heart, five isoforms predominate, including two long membrane-bound forms (L-OPA1) that are further cleaved into three soluble short forms (S-OPA1) (Garcia et al., 2021). A balanced L-/S-OPA1 ratio is essential for efficient inner mitochondrial membrane fusion. L-OPA1 contains two key cleavage sites: S1, targeted by the stress-activated protease OMA1, and S2, processed by the ATP-dependent protease YME1L (Zhang et al., 2014, Del Dotto et al., 2018).

Davies and his group (2007) found that homozygous inactivation of Opa1 leads to early embryonic lethality (Davies et al., 2007) in mouse. Studies on human cell lines revealed loss of OPA1 triggers the fragmentation of the Mitochondrial-network, disorganisation of cristae structure, dissipation of mitochondria membrane potential, reduced respiratory capacity, opening of cristae junction allowing mobilization of cytochrome c into the cytoplasm, which subsequently activates caspase dependent apoptosis (Olichon et al., 2003; Chen et al., 2005). OPA1 haploinsufficiency is responsible for the most common form of autosomal dominant optic atrophy (ADOA), an inherited optic neuropathy characterized by selective degeneration of retinal ganglion cells, resulting in optic nerve degeneration and visual failure (Votruba, 1998, Delettre et al.,2000; Del Dotto et al., 2018). Beyond isolated optic atrophy, pathogenic OPA1 variants have been also associated with spectrum of neurodegenerative phenotypes, such as Behr-like syndrome, syndromic parkinsonism and dementia (Marelli et al., 2011, Carelli et al., 2015; Del Dotto., 2018). More over OPA1 deficiency has been associated with cardiomyopathy, metabolic stoke and other inherited conditions (Chen et al., 2012; Zerem et al., 2019).

Opa1 plays a prominent role in cardiac development and function. In *Drosophila melanogaster*, heterozygous mutation of dOpa1 leads to decreased heart rate, increased heart arrhythmia, impaired fractional shortening and heightened susceptibility to heart failure in response to electrical pacing (Shahrestani et al., 2009; Dorn et al., 2011). Consistently, mouse model having reduced cardiac OPA1 expression exhibit profound mitochondria network remodelling, impaired direct channelling of ATP and ADP between mitochondria and myosin ATPases, more accumulation of calcium and delay in calcium-induced Mitochondrial Permeability Transition Pore (mPTP) opening and mice are more sensitive to prolonged haemodynamic stress (Piquereau et al., 2012).

In order to assess the involvement of OPA1 in the development of DCM, here we have investigated an uncommon missense variation in OPA1 (p.P188L), identified by whole-exome sequencing. *In vitro* functional analyses demonstrated a marked decrease in mitochondrial membrane potential and ATP synthesis, alongside heightened cytosolic calcium concentrations, augmented reactive oxygen species (ROS) production, and pronounced mitochondrial fragmentation in cells harbouring the p.P188L variant compared to wild-type. OPA1 Structural modelling forecasted significant alterations in the secondary and tertiary structures of the mutant protein as well as weakened binding with OMA1. These data collectively suggest that the OPA1 p.P188L variation disrupts mitochondrial dynamics and bioenergetics balance, which are critical aspects of mitochondrial dysfunction, therefore leading to compromised cardiac muscle performance. This study provides the inaugural evidence connecting an OPA1 variation to DCM within the Indian population.

## Material & Methods

### 2.1 Enrolments of study subject and clinical assessment

The current investigation included 5 families (FDCM1-FDCM5), each consisting of at least of two affected individuals and one unaffected member, in addition to 10 sporadic DCM cases (SDCM1-SDCM10), all of whom underwent whole exome sequencing (WES). Additionally, 100 age-matched healthy individuals without clinical indications of cardiac problems were included. The diagnosis was determined by left ventricular (LV) dimensions and functions, notably a Left Ventricular Ejection Fraction (LVEF) of less than 45% and an LV fractional shortening (LVFS) of less than 25%. Individuals with hypertension, coronary artery disease (CAD), or congenital heart disease (CHD) were excluded. The research obtained authorisation from the Institutional Ethics Committee.

### 2.2 Genomic DNA Extraction and Whole-Exome Sequencing

Whole-exome sequencing (WES) was conducted on trios, comprising the proband and first-degree relatives. Genomic DNA was extracted from peripheral venus blood samples via the standard ethanol precipitation method. The purity and concentration of DNA were checked employing NanoDrop 2000 spectrophotometer (Thermo Fisher Scientific). The probands from different families were chosen for WES, along with at least one affected and one unaffected family member (who acted as an internal control). Exome library preparation was conducted utilising the Human Nextera Rapid Capture Expanded Exome Kit (Illumina, USA) following the manufacturer’s guidelines. The generated libraries were sequenced using the Illumina HiSeq 2500 platform (Illumina, USA), producing 150 bp paired-end reads, with an average sequencing depth of around 100× per sample. The sequencing data underwent thorough quality control evaluations to assure integrity and dependability.

### 2.3 NGS data analysis

The data analysis was conducted subsequent to the completion of sequencing. The sequencing data were aligned to the human reference genome (GRCh38 or hg19) utilising alignment methods. Base quality score recalibration was executed utilising the ‘GATK Recalibrator’ to improve variant calling precision. Ultimately, variant calling was performed to provide VCF files for each sample. A comprehensive outline of the NGS data analysis workflow can be provided upon request.

### 2.4 Prioritization of Putative Pathogenic Variants

A panel of genes, encompassing those previously linked to heart development and disease, was assembled by a comprehensive literature review for curation of inhouse cardiac gene pipeline (∼3000 genes). The potentially harmful variations were prioritised utilising the pipeline. The variants having a read depth of 20 or greater were selected to prevent the omission of potential variations with insufficient read depth. The curated cardiac gene panel was employed to select the variations. Variants with a minor allele frequency (MAF) ≥ 0.01 in the Genome Aggregation Database (gnomAD; https://gnomad.broadinstitute.org/), 1000 Genomes Project (https://www.internationalgenome.org/), Exome Aggregation Consortium (ExAC) allele frequency database (http://exac.broadinstitute.org/), IndiGenomes database (https://clingen.igib.res.in/indigen/), and INDEX database (https://indexdb.ncbs.res.in/search/) were excluded. Subsequently, the selected variants were classified as missense, stop codon, frameshift, splice-site, synonymous, and intronic variants based on their impact on the protein sequence.

### 2.5 Variant confirmation and segregation Analysis

The selected variants were further confirmed by Sanger sequencing method using BigDye® Terminator v3.1 Cycle Sequencing Kit (Applied Biosystems, Inc.). The sequencing was performed on DNA Analyzer (ABI-3130, Applied Biosystems, USA). Sequencing data were analyzed by using FinchTV chromatogram viewer (http://www.geospiza.com/ftvdlinfo.html, Geospiza, Seattle, WA, USA). The segregation pattern of the identified variants were also performed in other family members of the proband by analysing their DNA sequence by Sanger sequencing method.

### 2.6 *In-silico* Prediction of Variant Pathogenicity

The pathogenic potential of non-synonymous variants in OPA1 was evaluated using VarCards (http://www.genemed.tech/varcards/search), which integrates over 20 plus *in silico* predictive algorithms, including PolyPhen-2, PANTHER, SIFT, Mutation Taster, I-Mutant, FATHMM, VEST3, M-CAP, PROVEAN, CADD and SNAP2 which collectively yield a damaging score ranging from 0 to 1, referred to as the VarCards score. Variants with a damaging score of 1 are classified as highly detrimental, whilst those with a value of 0 are deemed non-damaging.

### 2.7 Evolutionary Conservation Analysis of Identified OPA1 Variant

The evolutionary conservation of the identified OPA1 variant was examined at both genomic and proteomic levels. Nucleotide-level conservation was evaluated employing phylogenetic constraint-based computational techniques, such as GERP, phyloP, phastCons, and SiPhy, which estimate evolutionary constraints across several taxa. Simultaneously, protein-level conservation was assessed using cross-species comparison of amino acid sequences utilising the NCBI HomoloGene database (http://www.ncbi.nlm.nih.gov/homologene).

### 2.8 Prediction of Secondary and Tertiary Protein Structures

The secondary and tertiary structures of a protein are dictated by the sequence and physicochemical characteristics of its component amino acid residues. Non-synonymous variations can modify protein conformation based on the characteristics of the substituted side chains. The secondary structural characteristics of the wild-type and mutant OPA1 proteins were predicted utilising the PSIPRED bioinformatics program (http://bioinf.cs.ucl.ac.uk/psipred/). Simultaneously, three-dimensional homology models of the wild-type (WT) and mutant (MUT) OPA1 proteins were constructed by using i-TASSER. Structural models with confidence values beyond 90% were chosen for additional analysis. The resultant WT and mutant models were later superimposed utilising UCSF Chimaera, and structural discrepancies between WT and MUT-OPA1 were assessed by computing root-mean-square deviation (RMSD) values.

### 2.9 Prediction of protein stability

Various computational methods were utilised to estimate the impact of sequence alterations on the structural stability and dynamic properties of the mutant OPA1 protein. DynaMut2 (https://biosig.lab.uq.edu.au/dynamut2/) was utilised to forecast alterations in protein stability and flexibility generated by mutations by the integration of vibrational entropy and dynamic studies. MUpro (https://mupro.proteomics.ics.uci.edu/) use machine learning techniques to assess changes in protein stability owing to single-residue substitutions. CUPSAT (https://cupsat.brenda-enzymes.org/) forecasts the impact of amino acid substitutions on protein stability by employing environment-specific atomic potentials and torsion angle distributions. DUET (https://biosig.lab.uq.edu.au/duet/stability), which integrates the complementing mCSM and SDM methodologies, was employed to augment the accuracy of stability predictions. I-Mutant (https://folding.biofold.org/i-mutant/i-mutant2.0.html) offers assessments of stability alterations linked to single-point mutations, utilising both sequence- and structure-based methodologies. These methods collectively provide complementary insights into the molecular pathways via which variations affect OPA1 protein structure and function.

### 2.10 Protein-protein docking

To elucidate the molecular interaction between OPA1 and OMA1 molecular docking was conducted using the HADDOCK2.4 web server (Dominguez et al., 2003). The PDB structures of both interacting partners were uploaded to the server, and active (binding) residues were defined based on previously reported literature (Baker et al., 2014; Tobacyk et al., 2019; Alavi, 2021). The resulting docked complexes were subsequently analyzed and visually inspected using PyMOL (version 4.6) to assess interfacial contacts, and structural perturbations induced by the mutations.

### 2.11 *In Vitro* Functional Assessment of OPA1 Variant

#### 2.11.1 Molecular Cloning and Site-Directed Mutagenesis

The full length human OPA1 clone was kindly provided by Prof. Daniel Linseman (University of Denver, Denver, CO) and then subcloned into the mammalian expression vector pcDNA3.1/NT-GFP-TOPO™ (Thermo Fisher Scientific) following the manufacturer’s instructions. The OPA1 construct’s integrity was verified using Sanger sequencing. The wild-type (WT) OPA1 plasmid served as a template for the generation of mutant OPA1 (MUT- OPA1) constructs using site-directed mutagenesis, employing the QuikChange II XL Site-Directed Mutagenesis Kit (Agilent Technologies, Santa Clara, CA, USA) in accordance with the manufacturer’s guidelines. The successful integration of the intended mutation was verified by sequencing.

#### 2.11.2 Cell Culture and Plasmid Transfection

The H9c2 cell line (rat cardiomyoblast), sourced from the ‘National Centre for Cell Science’ (NCCS) in India, was utilized for the establishment of stable cell lines. Cells were cultured in Dulbecco’s Modified Eagle’s Medium (DMEM; Gibco, Life Technologies Corp.) enriched with 10% fetal bovine serum (FBS; Gibco) at 37 °C in a humidified atmosphere containing 5% CO . Cells were inoculated in 6-well tissue culture plates at about 40-45% confluency and transfected with the OPA1-WT expression plasmid, 24 hours post-seeding with FuGENE®-6 HD transfection reagent (Promega Corp.), in accordance with the manufacturer’s guidelines. Post-transfection, cells were permitted to recuperate for 48 hours in full growth media in order to ensure adequate transgene expression. A G-418 disulfate kill curve was established to ascertain the minimum antibiotic concentration necessary to eradicate non-transfected cells. Selection was conducted utilizing the optimal G-418 concentration, ensuring that only cells with stably integrated plasmids expressing the neomycin resistance gene survived, while untransfected cells were eliminated. Upon the visibility of antibiotic-resistant colonies, individual clones were separated using limiting dilution and subsequently grown in DMEM supplemented with 10% FBS and G-418. Subsequent confirmation of stable integration and expression was achieved using quantitative real-time PCR. Following the validation of the OPA1-WT stable cell line, similar approach was utilized to produce OPA1-P188L mutant stable cell line.

#### 2.11.3 OPA1 Expression in Cultured Myoblasts

To investigate the cellular distribution of wild-type and mutant OPA1 variants, stable H9c2 cell lines harbouring these constructs were cultured on glass coverslips situated in 6-well plates. Cultures were cultivated overnight at 37°C in a 5% CO atmosphere. Upon achieving 70-80% confluency, the cells underwent two washes with 1X PBS, then treated with 50nM MitoTracker Red CMXROS (Molecular Probes) for 30 minutes at 37 C, and then fixed using 4% paraformaldehyde (PFA) for 10 minutes. Before immuno-staining, the cells were made permeable using 0.1% Triton X-100 diluted in 1X PBS for 15 minutes, then incubated for 2 hours in blocking buffer (1% BSA in PBS) at room temperature. The cells were then treated overnight at 4°C with a primary antibody (Anti-OPA1 antibody, Cell Signalling Technology, USA, diluted 1:100) in 0.1% BSA. Following extensive rinsing with 1X PBST (5X, 5 minutes each), the specimens were incubated with Alexa Fluor 488-conjugated Goat anti-rabbit IgG (Molecular Probes, OR, USA) for 2 hours at room temperature. Following incubation, the cells underwent thorough washing with 1X PBST (5-6 times), succeeded by nuclear staining utilising DAPI. The stained cells were affixed with mounting media and visualised with the SP8 STED Laser Scanning Super-Resolution Microscope System from Leica. Image analysis was conducted utilising Image J software (NIH, Bethesda, MD, USA).

#### 2.11.4 Western blot analysis of protein expression

The impact of OPA1 missense mutations on protein expression was estimated by conducting ‘Western blot’ analysis. Stable OPA1-expressing cell lines, comprising wild-type (OPA1- WT) and mutant OPA1- MUT (P188L), were plated at equal densities (10^5^/well) in six-well plates and collected upon achieving above 90% confluency. Whole-cell lysates were generated utilizing RIPA buffer (50 mM Tris–HCl, pH 8.0; 150 mM NaCl; 0.1% Triton X- 100; 0.5% sodium deoxycholate; 0.1% SDS) augmented with 1 mM sodium orthovanadate, 1 mM sodium fluoride, and a 1× protease inhibitor cocktail (Roche, Switzerland). Proteins were separated using 10% SDS–polyacrylamide gels and subsequently transferred to PVDF membranes (Bio-Rad Laboratories). Membrane was incubated for 2 hours at room temperature with 5% (w/v) non-fat dry milk in TBST (0.1% Tween-20) to block, followed by overnight incubation at 4 °C with an OPA1-specific primary antibody (Cell Signalling Technology, USA; 1:1000). After the primary antibody incubation, membranes were washed thrice with 1XTBST (5 minutes each) and subsequently incubated for one hour at room temperature with HRP-conjugated goat anti-mouse IgG secondary antibody (Genei, Merck Specialties Pvt. Ltd.). Following five additional washes with TBST (5 minutes each), immunoreactive bands were detected utilizing an enhanced chemiluminescence (ECL) detection technique (GE Healthcare, IL, US).

#### 2.11.5 MTT-Based Cell Viability Assay

Stable OPA1 wild-type (WT) and mutant (MUT) cell lines were inoculated onto 96-well plates in triplicate, roughly at a density of 2 × 10 cells per well. After 24 hours, MTT reagent was introduced to each well at a final concentration of 0.5 mg/mL, followed by incubation for 4 to 5 hours at 37 °C in a humidified environment with 5% CO . The resultant blue formazan crystals were solubilized in dimethyl sulfoxide (DMSO) and incubated in darkness for 10 minutes. The formation of formazan indicates mitochondrial metabolic activity and acts as an indirect indicator of cell viability (Riss et al., 2016). Absorbance was quantified at 570 nm with a microplate reader (Bio-Rad), and cell viability was determined in accordance with the standard formula.

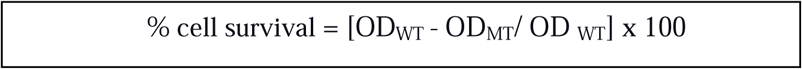

#### 2.11.6 Measurement of Mitochondrial Membrane Potential (**ΔΨ**m)

Stable cell lines expressing OPA1-WT and OPA1- MUT constructs were maintained in 6- well plates. After 24 hours, cells were rinsed with phosphate-buffered saline (PBS), trypsinized, and gathered in 1.5 mL microcentrifuge tubes. Cells were subsequently stained with 200 nM tetramethyl-rhodamine ethyl ester (TMRE; Invitrogen) and incubated for 30 minutes at 37 °C in the dark. The mitochondrial membrane potential-dependent TMRE fluorescence was measured via flow cytometry utilizing a ‘Cyto-FLEX LX’ system (Beckman Coulter). The fluorescence intensity of cells positive for the orange channel was quantified as a measure of mitochondrial membrane potential. Carbonyl cyanide-p-trifluoromethoxy-phenyl-hydrazone (FCCP; 10 μM, 10 min) used as a positive control to cause whole mitochondrial membrane depolarization and to confirm the specificity of TMRE fluorescence for ΔΨm. The data are expressed as a geometric mean fluorescence intensity (gMFI) ± Standard error of mean (SEM) from 6 independent experiments.

#### 2.11.7 Quantification of Cellular ATP Content

Stable WT and mutant OPA1-expressing cell lines were inoculated in 12-well culture plates at uniform densities (∼10²· cells per well) and incubated for 24 hours. Subsequent to incubation, cells were rinsed twice with 1× phosphate-buffered saline (PBS), and intracellular ATP concentrations were measured utilizing the ENLIGHTEN ‘ATP Assay’ System (Promega, WI, USA) in accordance with the manufacturer’s instructions. Before sample analysis, a standard curve was established with known ATP values to enable accurate estimation. Cellular ATP was extracted using 0.5% trichloroacetic acid (TCA) and subsequently neutralized with 0.1 M Tris–acetate buffer at pH 7.75. Equal amounts of recombinant luciferase/luciferin (rL/L) reagent was used, and luminescence was measured with a luminometer at two intervals (2 seconds and 10 seconds). The cellular ATP content was determined as the average of the luminescence values recorded at both time intervals. The data are expressed as a geometric mean luminescence intensity (gMFI) ± Standard error of mean (SEM) from 6 independent experiments.

#### 2.11.8 Assessment of Mitochondrial Mass Using NAO

Stable OPA1-WT and mutant OPA-MUT cell lines were plated in 6-well plates at uniform densities (1 × 10 cells per well) and cultured for 24 hours. Subsequently, cells were trypsinized and resuspended in fresh complete medium containing 1 μM 10-N-nonyl acridine orange (NAO; Invitrogen), followed by incubation for 30 minutes at 37 °C in a humidified environment with 5% CO, as previously documented (Maftah et al., 1989). After incubation, cells were washed with 1× phosphate-buffered saline (PBS) and resuspended in PBS enriched with calcium and magnesium ions. Fluorescence measurements were obtained using a ‘FACSCalibur’ flow cytometer at an excitation/emission wavelength of 495/522 nm. For each sample, 10,000 events were recorded. The data are expressed as a geometric mean fluorescence intensity (gMFI) ± Standard error of mean (SEM) from 6 independent experiments.

#### 2.11.9 Flow Cytometric analysis of Oxidative Stress

Stable WT and MUT-OPA1-expressing cell lines were seeded in 6-well plates at a density of 1 × 10 cells per well and incubated for 24 hours. Cells were then trypsinized and resuspended in fresh complete media containing MitoSOX™ Green (5 μM), followed by incubation for 10 minutes at 37 °C. After staining, fluorescence was quantified using a FACSCalibur flow cytometer (Bio-Rad) (Winyard et al., 2011; Jiang et al., 2016). The data are expressed as a geometric mean fluorescence intensity (gMFI) ± Standard error of mean (SEM) from 6 independent experiments.

#### 2.11.10 Measurement of Intracellular Ca² Levels

OPA1 stable cell lines harboring wild-type (WT) and mutant (MUT) constructs were inoculated in 6-well plates at a density of 1 × 10 cells per well and maintained under standard growth conditions for 24 hours. Cells were subsequently washed with phosphate-buffered saline (PBS), trypsinized, and resuspended in fresh culture media containing Fluo-3 AM (Invitrogen) at a final concentration of 0.5 μM. The cell suspensions were incubated for 30 minutes at 37 °C in the absence of light to facilitate optimal dye loading. After incubation, fluorescence was quantified with a FACSCalibur flow cytometer (Bio-Rad). The data are expressed as a geometric mean fluorescence intensity (gMFI) ± Standard error of mean (SEM) from 6 independent experiments.

#### 2.11.11 Quantitative Analysis of Mitochondrial Oxygen Consumption Rate (OCR)

Mitochondrial respiration was measured using a high-resolution respirometer (Oxygraph-2 k, OroborosInstruments) at 37°C under stirring conditions (750 rpm). Stable cell lines of OPA1- WT and OPA1-MUT cells were trypsinized, and washed twice and resuspended mitochondria respiration buffer. A total of 1X10^6^ cells were introduced into each oxygraph chamber, with separate chamber containing OPA1-WT and OPA1-MUT. Respiration was allowed to determine routine respiration, reflecting the physiological coupling state under endogenous substrate supply and ATP demand. Then, Cells were then sequentially treated with oligomycin (1 μg/ml), FCCP (0.5 µM), and antimycin A (1 μg/ml) to study leak respiration, maximum capacity of electron transport system and residual/extra-mitochondrial oxygen consumption, respectively. Calibration at air saturation was performed each day before starting experiments by letting Buffer B stir with air in the oxygraph chamber until equilibration and a stable signal was obtained. All experiments were performed at an oxygen concentration in the range of 100–205 μM O2. Data were recorded and analyzed using DatLab 7.4 software (Oroboros Instruments).

#### 2.11.12 Quantitative PCR Analysis of Gene Expression

Quantitative real-time PCR (qRT-PCR) was conducted to assess the mRNA expression levels of mitochondrial DNA-encoded respiratory chain subunits (MT-ND1 and MT-CO1) and apoptosis-related genes (Bax and Bcl-2) in stably transfected H9c2 cell lines expressing OPA1-WT and OPA1-MUT. Total RNA was extracted from the stable cell lines utilizing the TRIzol reagent in accordance with the manufacturer’s guidelines and subsequently subjected to treatment with RNase-free DNase I (Thermo Fisher Scientific, USA). RNA concentration was quantified using a NanoDrop 2000 spectrophotometer (Thermo Fisher Scientific), and RNA purity was evaluated by the A260/A280 ratio, RNA integrity was verified by electrophoresis on a 1% (w/v) agarose gel (A9539, Sigma-Aldrich). Complementary DNA (cDNA) was generated from total RNA utilizing the RevertAid First Strand cDNA Synthesis Kit (Thermo Fisher Scientific). Real-time PCR was conducted with the KAPA SYBR® FAST qPCR Master Mix Kit (Merck KGaA, Darmstadt, Germany) on a QuantStudio™ 5 Real-Time PCR System (Applied Biosystems, USA). β-Actin served as the endogenous internal control for normalization purposes. All experiments were conducted in triplicate, quantification was expressed as fold change with the standard error of mean and the p value when <0.05 was considered statistically significant.

#### 2.11.13 Prediction of RNA secondary structure

After evaluating protein instability, we broadened our investigation to examine mRNA architecture at the transcript level. To do this, the desired RNA sequence was derived from the OPA1 transcript (NM_015560). To achieve this objective, small RNA fragments were created, each comprising 41 nucleotides in total. To precisely evaluate the effect of the alteration, the mutation was deliberately located at the 21st nucleotide, thus centring it within a segment that encompassed 20 nucleotides on both sides, offering an extensive context for the sequence milieu surrounding the modification. After the RNA sequences were generated, the secondary structures of both wild-type and mutant RNA segments of OPA1 were examined utilising the ‘RNAStructure’ Web Server (version 6.0.1) (Bellaousov et al., 2013). This platform employs thermodynamic algorithms to forecast RNA secondary structures, offering detailed understanding of folding configurations and base-pairing interactions in RNA sequences. Furthermore, the ’MutaRNA’ tool was utilised to conduct a mutational study of RNA segments, investigating the structural alterations caused by each missense variant (Miladi et al., 2020).

#### 2.11.14 Statistical analysis

Data obtained from Western blot and RT-qPCR experiments were expressed as mean fold change ± Standard error of mean (SEM) from three independent experiments. Statistical analyses were conducted using unpaired Student’s t-test to compare differences between WT and experimental groups individually. A p-value < 0.05 was considered statistically significant. Similarly, statistical significance of all other experments including measurement of ATP content, mitochondrial membrane potential, OCR, mitochondrial mass, Ca2+ homeostasis, ROS production and mtDNA content was calculated using unpaired Student’s t-test.

## 3. Results

### 3.1 Identification of OPA1 rare variant associated with DCM

Whole-exome sequencing of five families and in ten isolated cases of dilated cardiomyopathy (DCM), discovered a solitary missense variant in the OPA1 gene, c.563C>T (p.Pro188Leu), with a minor allele frequency of 0.000069 in the gnomAD database (rs772090345). The proband was a 5-month-old male exhibiting significantly impaired cardiac function, characterized by a markedly diminished left ventricular ejection fraction (LVEF = 21%) and an enlarged left ventricular internal diameter in diastole (LVIDd = 89 mm) (Figure 2C), indicative of severe left ventricular systolic dysfunction. The mutation was then confirmed using PCR amplification and Sanger sequencing in the proband (Figure 2D) and available family members. The mutation was identified alone in the proband and was not present in any other family members, indicating its *de novo* emergence.

**Figure 1:**
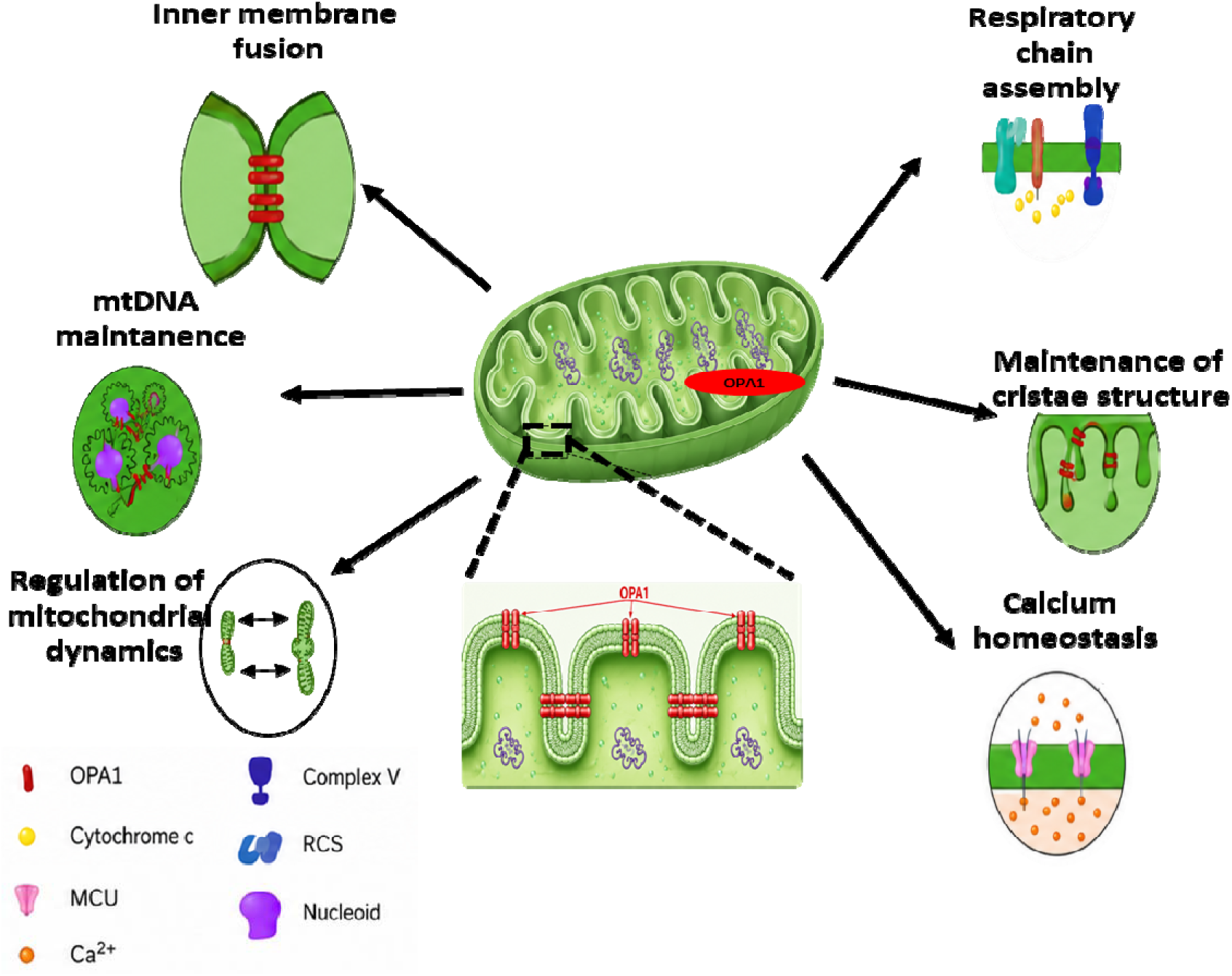
Schematic illustration of OPA1-mediated regulation of mitochondrial fusion, cristae architecture, calcium signaling, and respiratory chain system.

**Figure 2:**
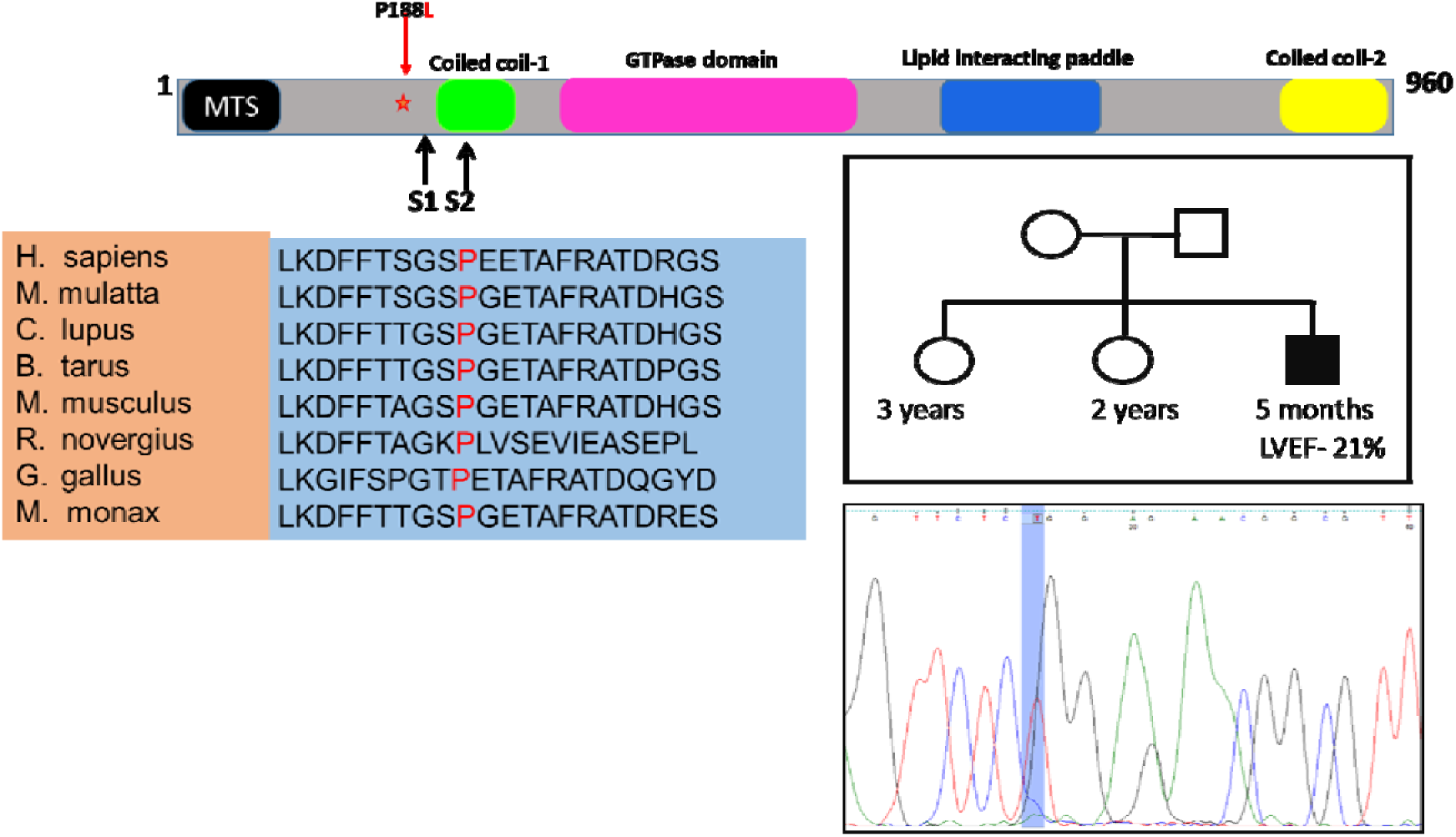
Schematic representation of localization, sequence chromatograms, pedigree and phylogenetic conservation of OPA1 non-synonymous variants. **(A)** OPA1 protein showing a MTS (Mitochondrial targeting sequence), Coiled coil domain1, GTPase domain, Lipid-interacting paddle/LIS and Coiled coil domain2. S1 and S2 are the site for proteolytic cleavage. Position of identified non-synonymous variants (present study) are marked in red colour **(B)** Phylogenetic conservation of OPA1 protein across different species showing substituted amino acid residues highlighted in different colours **(C)** Pedigree of the patient carrying the p.P188L OPA1 variant. Circles correspond to females, squares to males. The black symbol indicates the proband affected by DCM, whereas the white ones are used for healthy relatives **(D)** Sequence chromatogram of nonsynonymous variant c.563C>T.

### 3.2 Cross-Species allignment of OPA1 protein

A cross species multiple sequence alignment of OPA1 protein across several species viz., *Homo sapiens* (NP_056375.2), *Macaca mulatta* (XP_014988011.1), *Canis lupus familiaris* (XP_022264504.1), *Bos taurus* (XP_005201558.1), *Mus musculus* (NP_598513.1), *Rattus norvegicus* (NP_001420836.1), *Gallus gallus* (NP_001383101.1), *Marmota monax* (NW_026693473.1) revealed that the substituted amino acid residue P188 was highly conserved. The conserved region showing the substituted amino acid along with flanking amino acid sequences has been shown in Figure 2B.

### 3.3 Alterations in protein stability

While evaluating the influence of the OPA1- P188L mutations on protein stability, using a range of *in-silico* prediction tools, it was demonstrated that the p.P188L substitution sustained high degree of structural destabilization, as consistently indicated across multiple protein stability prediction platforms (Table 2)

**Table 1.** In silico analysis of identified OPA1 variant across different population database.

| Nucleotide change | Amino Acid change | Domain | dbSNP ID | Type of mutation | Status | ClinVar Annotation | InterVar | Genome Asia 100k | HGMD Disease | 1000 Genome | genomeAD |
| --- | --- | --- | --- | --- | --- | --- | --- | --- | --- | --- | --- |
| c.563C>T | p.Pro188Leu | Linker | rs772090345 | Non-Synonymous | Known | NR | Uncertain | NR | NR | NR | 0.0000690675 |

**Abbreviations:** dbSNP: Single nucleotide polymorphism database, NR: Not reported

**Table 2.** Prediction of pathogenic likelihood of OPA1 variant using VarCards and structure stability assessment.

| Variant | Sift | Polyphen2 | LFT pred | Mutation tester | FATHMM | PROVEN | CADD | DYNAMUT2 | MUpro | CUPSAT | DUET | I-Mutant | SDM | mCSM |
| --- | --- | --- | --- | --- | --- | --- | --- | --- | --- | --- | --- | --- | --- | --- |
| c.563C>T | Damaging | Damaging | Damaging | Disease causing | Damaging | NR | Damaging | -0.89kcal/mol Destabilizing | -0.89kcal/mol Destabilizing | -2.17kcal/mol Destabilizing | -0.995kcal/mol Destabilizing | Decrease | -0.85kcal/mol Destabilizing | -0.495kcal/mol Destabilizing |

SIFT: sorting intolerant from tolerant; PolyPhen2: polymorphism phenotyping v2; FATHMM: functional analysis through hidden Markov models; CADD: combined annotation dependent depletion Dynamut2: dynamic Mutant 2; MUpro: mutant protein predictor; CUPSAT: Cologne University Protein Stability Analysis Tool; DUET: Double Mutation Estimator Tool; I-mutant: internet-based mutant; SDM: site-directed mutator; mCSM: mutation cutoff scanning matrix.

### 3.4 Predicted Conformational Alterations in OPA1 and Their Influence on OMA1 Binding

The three-dimensional structures of both the wild-type (WT) and mutant (MUT) forms of the OPA1 protein were modelled and subsequently superimposed to assess the structural deviations induced by the mutation. Comparative structural analysis revealed a notable conformational change between the two models, as reflected by a root mean square deviation (RMSD) value of 3.5 Å (figure 3), indicative of substantial atomic displacement. Such a high RMSD value suggests that the mutation induces pronounced conformational alterations, potentially disrupting the native folding pattern, domain orientation, and interaction interface of OPA1, which may in turn compromise its functional association with interacting partner proteins.

**Figure 3:**
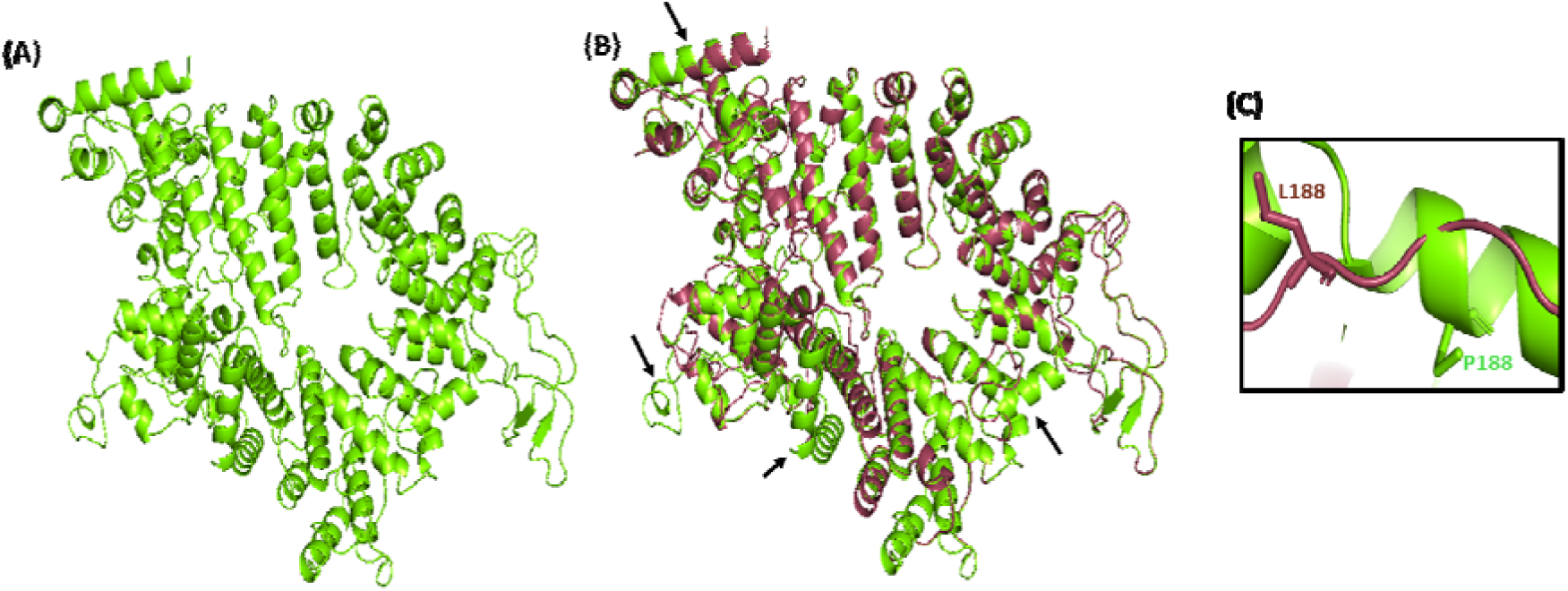
Structural comparison of OPA1-WT and OPA1-MUT variant by UCSF Chimera. **(A)** Three-dimensional structures of OPA1-WT. **(B)** Superimposed three-dimensional structures of wild type (OPA1-WT) versus mutant protein (OPA1-MUT) using Chimera. The root mean square deviation (RMSD) value of the two superimposed structures is 3.5. WT tertiary structure of MFN2 protein has been shown in green color while MUT has been shown in red color. **(C)** WT and MUT three-dimensional structure showing amino acid, Proline (P) and MUT amino acid, leucine (L) with neighboring amino acids.

For example, *In-silico* docking study of OPA1 and the mitochondrial protease OMA1 demonstrated significant disparities between the wild-type and P188L mutant complexes. The mutant OPA1 had an expanded interaction interface area and an increased number of interface residues relative to the wild-type complex. Nonetheless, the mutant complex exhibited the reduction in hydrogen bonds and non-bonded interactions. These data indicate that mutation-induced structural alterations enlarge the interaction surface and may enhance the accessibility of OMA1, which might be promoting OPA1 proteolytic processing and their by increased in number of fragmented mitochondria. The detailed docking conformations and interactions are illustrated in (Figure 4)

**Figure 4:**
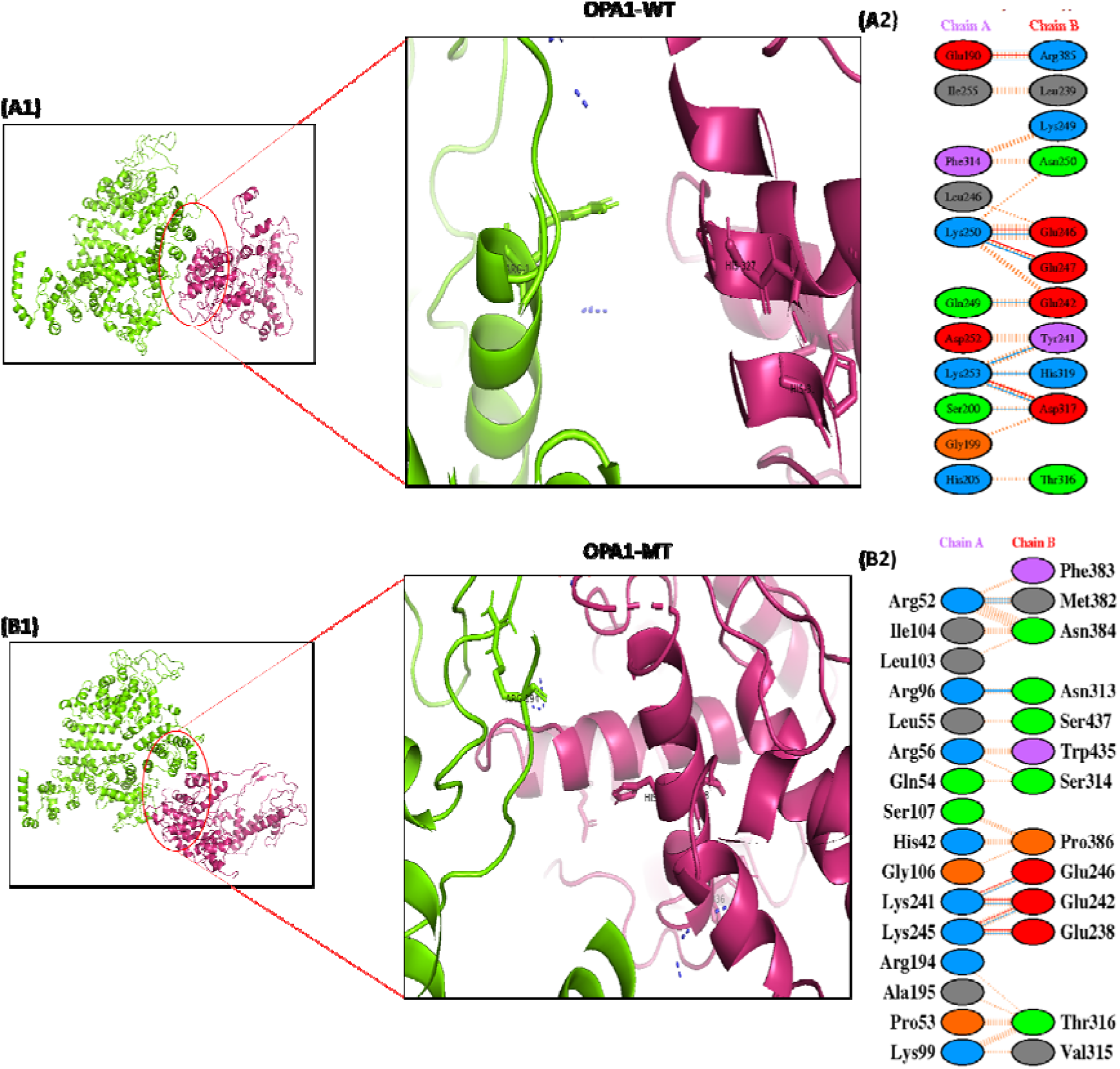
Protein-protein interaction of OPA1 protein with OMA1; **(A1,B1)** represents protein-protein interactions of Wild-type OPA1 (Green) as well as mutant OPA1(P188L) with OMA1 (Magenta). **(A2,B2**) represents residue involved in interaction.

### 3.5 Impact of P188L variant on the expression and localization of OPA1 protein

The expression of wild-type (WT) and mutant (P188L) OPA1 proteins in stably transfected H9C2 cells was assessed by Western blot analysis. One long (∼95 kDa) and a short (∼75 kDa) isoforms of OPA1 were detected as two separate immunoreactive bands. When compared to WT cells, both isoforms showed a notable decrease in the protein expression in cells bearing the p.P188L variation. The p. P188L variation significantly reduces OPA1 protein expression, as evidenced by the quantitative densitometric measurement of total OPA1 protein levels, which showed a substantial ∼1.7 fold down-regulation in OPA1-MUT cells compared to WT controls (p < 0.005) (Figure 5B).

**Figure 5:**
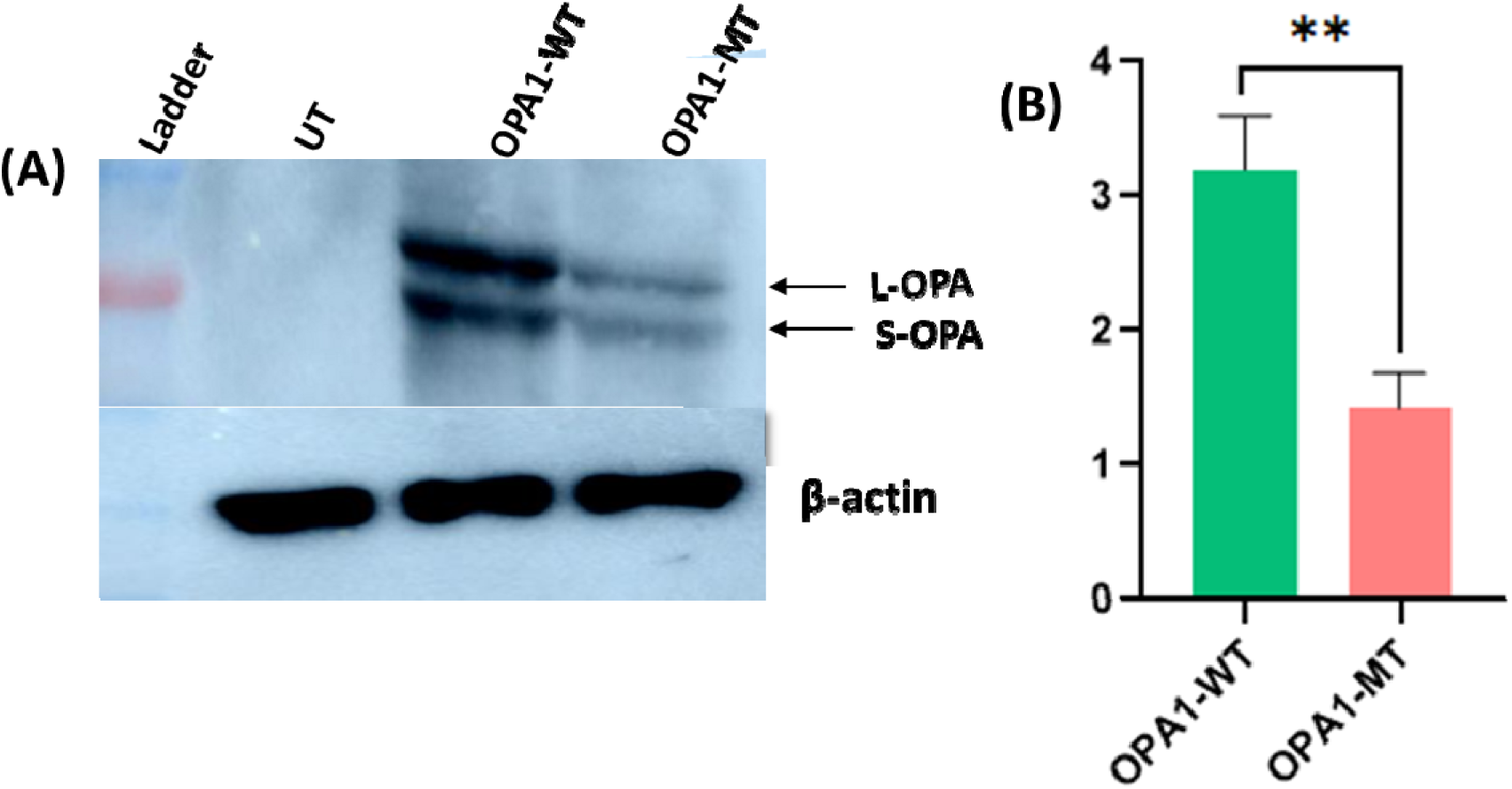
**(A)** Western blot analysis of wild-type and mutant protein showing expression of OPA1 **(B)** Representation of protein expression in terms of fold changes, calculated for each of the variant (P<0.05)

*In vitro* localization experiments in H9C2 cells revealed that both wild-type (WT) and mutant OPA1 proteins were mostly confined to the mitochondria. Cells harbouring the mutant OPA1 protein demonstrated a substantial decrease in OPA1 levels accompanied by pronounced mitochondrial morphological defects. Mito-Tracker labeling demonstrated that cells carrying the OPA1 p.P188L mutation exhibited significant mitochondrial fragmentation (Figure 6A). Quantitative analysis revealed that OPA1-MUT cells exhibited 67 ± 6.2% fragmented mitochondria, 12 ± 3.4% elongated tubular mitochondria, and 22 ± 5.4% mitochondria with intermediate morphology. In contrast, WT cells mostly displayed an interconnected mitochondrial network, comprising 24 ± 3.8% elongated tubular mitochondria, 22 ± 4.1% fragmented mitochondria, and 49 ± 5.4% intermediate mitochondria (Figure 6B). The observed mitochondrial fragmentation is likely a result of diminished OPA1 levels, which impairs inner mitochondrial membrane fusion and destabilizes cristae architecture, thereby leading to increased mitochondrial fragmentation.

**Figure 6:**
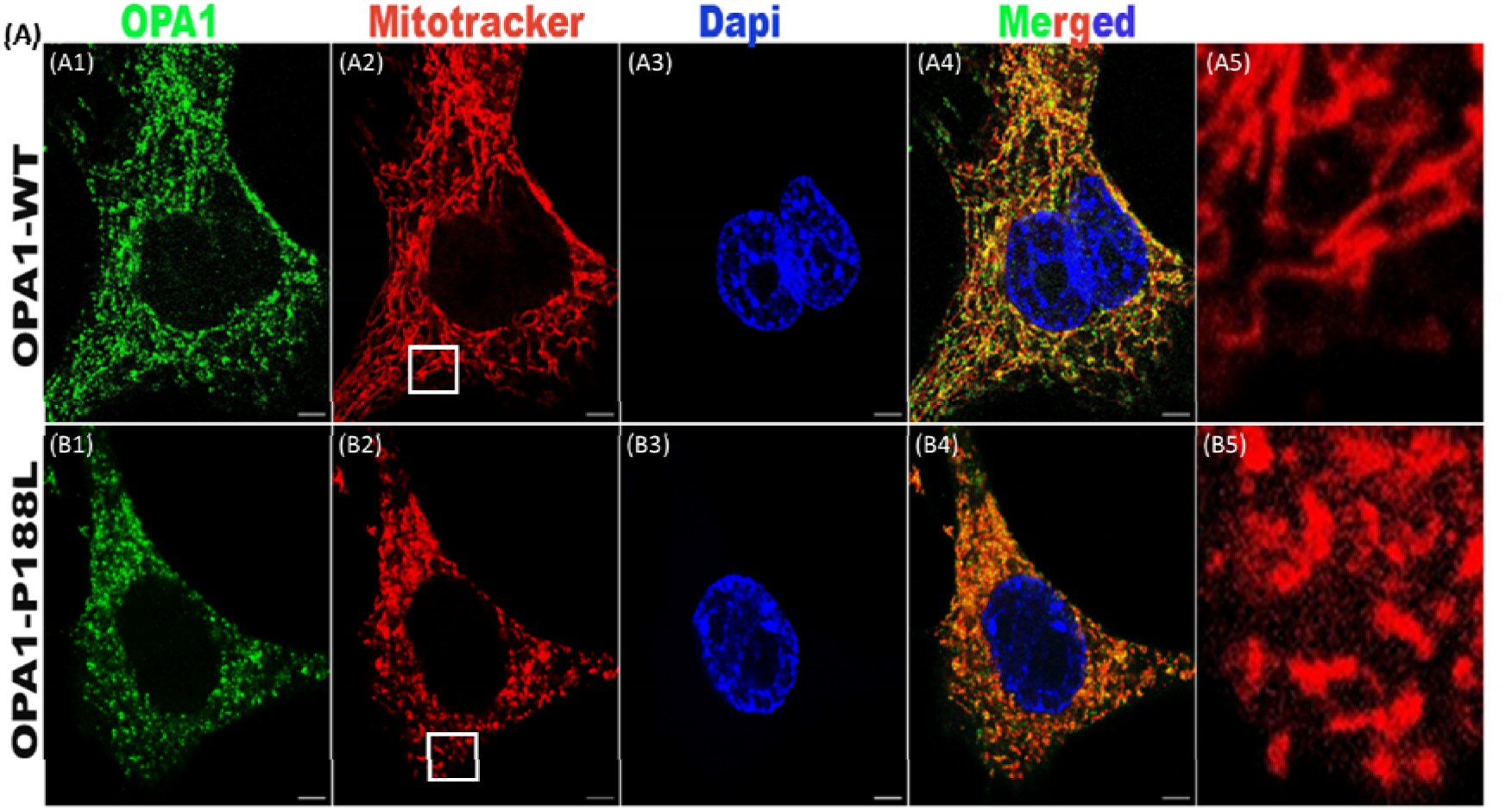
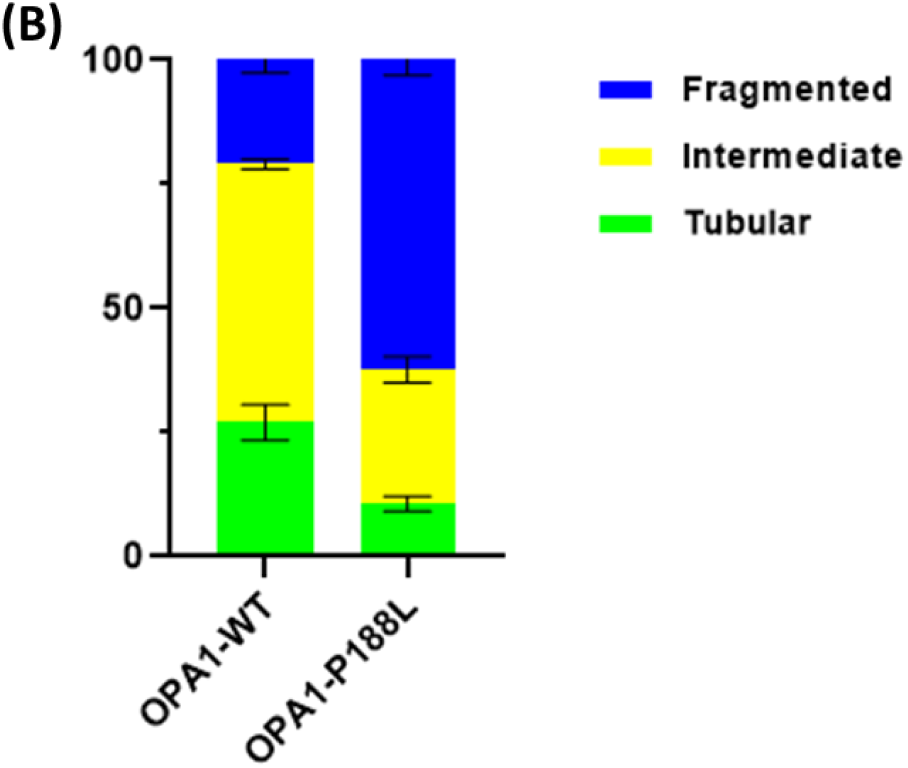
**A.** Expression analysis of OPA1 protein (A1-B4) Immunostaining in H9C2 cells showing cellular localization of OPA1-wild-type and mutant protein (p.P188L), A5 & B5 show higher magnification from boxed area. **B.** Quantification of mitochondria morphology in the cell lines described, Data are presented as mean ± SEM from three independent blind experiments, with approximately 100 cells analyzed per experiment.

### 3.6 Effect of P188L variant on cell viability and mitochondria membrane potential

To estimate the impact of the pP88L variant in the OPA1 gene on cellular health and mitochondrial function, a series of functional assays were performed. Cell viability was initially checked using the MTT assay. The results disclosed no significant difference in the percentage of viable cells between WT and MUT cells, suggesting that the P188L variant does not exert an immediate cytotoxic effect under the experimental conditions tested (Figure 7A). However, when we evaluated mitochondrial health by measuring mitochondrial membrane potential using TMRE (Tetramethyl rhodamine methyl esters) a fluorescent dye, used to measure (ΔΨm), we observed a (0.27 fold, ∼27%, p<0.01) notable decrease in mitochondrial memberane potential (ΔΨm) in OPA-MUT cells compared to WT cells (Figure 7B). This reduction in membrane potential is significant, as it can lead to impaired oxidative phosphorylation and subsequently causing significant decreased in ATP production (Figure 7C).

**Figure 7:**
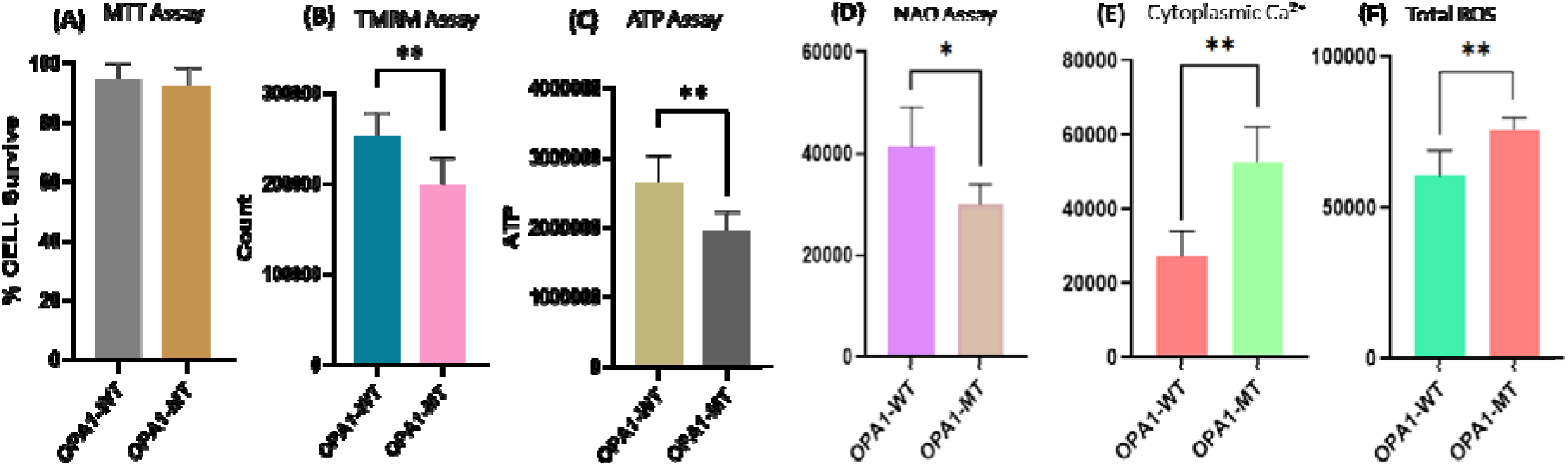
Effect of P188L on cell viability, mitochondria membrane potential, ATP level, mitochondrial mass, cytopasmic calcium level and total ROS. **(A)** Analysis of cell viabilty using MTT assay **(B)** Analysis of mitochondrial membrane potential: histogram shows fluorescent intensity values. **(C)** Luciferase assay to measure ATP level: histogram shows bioluminescence intensity-based total ATP level **(D)** Mitochondrial mass assay (NAO) **(E)** Cyotplasmic Ca² level measurement between OPA1-WT and P188L **(F)** Measurement of total ROS level. Mean values and standard errorS of mean were calculated from at least 6 independent experiments., *p < 0.05. **p<0.01 . (WT = wild-type)

### 3.7 Effect of OPA1-MUT on Mitochondrial Oxygen consumption rate and ATP content

Consistent with lowered mitochondrial membrane potential, the Oxygen consumption rate (OCR) when measured using a mitochondrial stress test, mitochondrial bioenergetics was found to be remarkably perturbed. Mitochondrial respiration was evaluated with the sequential administration of oligomycin (an ATP synthase inhibitor), FCCP (a protonophore and mitochondrial uncoupler), and the electron transport chain inhibitors rotenone and antimycin A. Respiratory metrics, including routine respiration, ATP-linked respiration (OXPHOS), maximal respiration, and residual oxygen consumption (ROX), were assessed. High-resolution respirometry analysis revealed a marked decline in mitochondrial respiratory function in OPA1-P188L cells relative to OPA1-WT controls. Routine respiration was markedly reduced (p < 0.01), signifying impaired basal mitochondrial activity. ATP-linked respiration (OXPHOS) was significantly decreased (p < 0.05), indicating compromised oxidative phosphorylation and lower ATP production capability. Maximal respiration rate was markedly diminished in mutant cells (p < 0.01), possibly indicating impaired electron transport chain capacity and reduced mitochondrial respiratory reserve. Moreover, residual oxygen consumption (ROX) was markedly reduced in OPA1-P188L cells (p < 0.01). Collectively, these results demonstrated a significant disruption in mitochondrial bioenergetics attributable to the P188L mutation (Figure 8A-B). Consistent with this, we also measured ATP concentration, which demonstrated a decrease in ATP levels (0.36 fold, ∼36%, p<0.01) in the P188L cells, suggesting that the P188L mutation in the OPA1 protein compromises mitochondrial integrity, potentially disrupting energy metabolism and cellular functions reliant on ATP.

**Figure 8.**
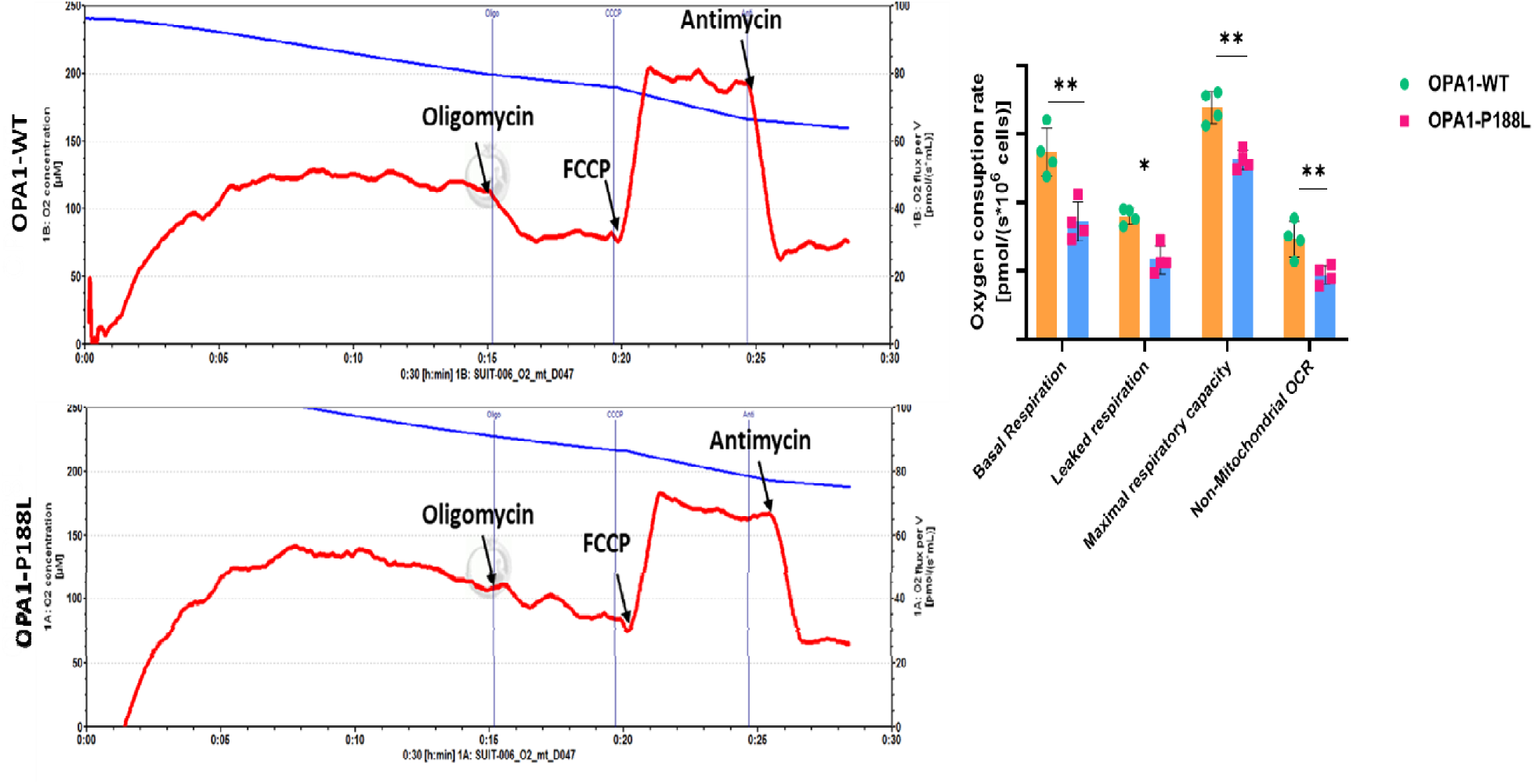
**(A & B)** Representative polarograms showing oxygen flux or oxygen concentration in H9C2 cells suspended in respiration buffer. Stable cell lines expressing OPA1-WT and OPA1-MUT were introduced into the oxygraph chamber. The blue tracing represents oxygen concentration within the chamber while the red trace indicates the rate of oxygen consumption (oxygen flux) by the cells. **(C)** Corresponding bar graph showing routine respiration, ATP-linked respiration (OXPHOS), leak respiration (LEAK) and residual oxygen consumption (ROX) rates in OPA1-WT and OPA1-P188L cells. Data are presented as mean ± standard error of mean. *p < 0.05 **p<0.01 (WT = wild-type).

### 3.8 Influence of OPA1-MUT on mitochondria mass

Further we have checked the mitochondria mass by using NAO (10-N-nonyl acridine orange), it binds with cardiolipin which is present in the inner membrane of mitochondria independent mitochondrial membrane potential (ΔΨm), serves as the primarily quantative marker of Mitochondrial mass, Integrity and organization of the inner mitochondrial membrane Cardiolipin content (Maftah et al., 1989; Petit et al., 1992), we have found reduced (0.35 fold, ∼35%, p<0.01) significant difference in MUT as compared to WT OPA1 (Figure 7D).

### 3.9 Calcium homeostasis and ROS generation

Immunostaining examination demonstrated a heightened quantity of fragmented mitochondria in the P188L-MUT cells. Biochemical studies consistently revealed a considerable disruption in mitochondrial membrane potential (ΔΨm) and diminished ATP generation, signifying impaired mitochondrial activity. These modifications indicate altered calcium homeostasis and oxidative stress. To explore this further, we measured total cytosolic calcium and reactive oxygen species (ROS) levels. Our findings demonstrated a substantial elevation in cytosolic Ca² (p < 0.01) and total ROS levels (∼ 24%, p < 0.01) in P188L-MUT cells relative to OPA1-WT cells (Figure 7E-F). The findings indicate that the P188L mutation modifies cristae junction architecture, resulting in diminished mitochondrial Ca² buffering capability and consequently increased cytosolic Ca² levels, corroborating earlier studies (Kushnareva et al., 2013).

### 3.10 OPA1 variant induce mtDNA depletion and activate apoptotic signalling in H9C2 cardiomyocytes

Mutations in OPA1 were anticipated to impact mitochondrial DNA (mtDNA) copy number due to alterations in maintenance of mitochondrial genome (Elachouri et al., 2011). To further examine this, mtDNA content was determined via real-time PCR employing the mitochondrial genes MT-ND1 and MT-CO1, with the nuclear gene β-actin utilised for normalisation. Our findings demonstrated a significant decrease in mtDNA copy number in OPA1-mutant cells, evidenced by a significant reduction in MT-ND1 (0.69-fold, p < 0.01) and MT-CO1 (0.70-fold, p < 0.01) compared with OPA1-WT cells. Quantitative RT-PCR analysis revealed altered expression of the mitochondrial-encoded genes *MT-ND1* and *MT- CO1* in OPA1-mutant cells, indicating compromised mitochondrial transcription, respiratory chain integrity and activation of intrinsic apoptotic pathways through cytochrome c release and caspase activation (Wang, 2001). Hence we further checked the expression of Bax, Bcl2, Caspase3 and 9 transcript levels and calculated the Bax/Bcl-2 ratio. A statistically significant increase in apoptotic index was observed in the P188L variant, in expression of caspase 9 (1.65-fold change, P < 0.051), caspase 3 (1.21-fold change, P < 0.01) and Bax/Bcl2 (1.29- fold change, P < 0.05), indicating a subtle shift toward apoptotic susceptibility (Figure 9A-F). Collectively, these findings demonstrate that OPA1 pathogenic variant induces mitochondrial dysfunction and apoptotic priming without promoting full apoptotic execution, reflecting their potential contribution to cardiomyopathy remodelling

**Figure 9:**
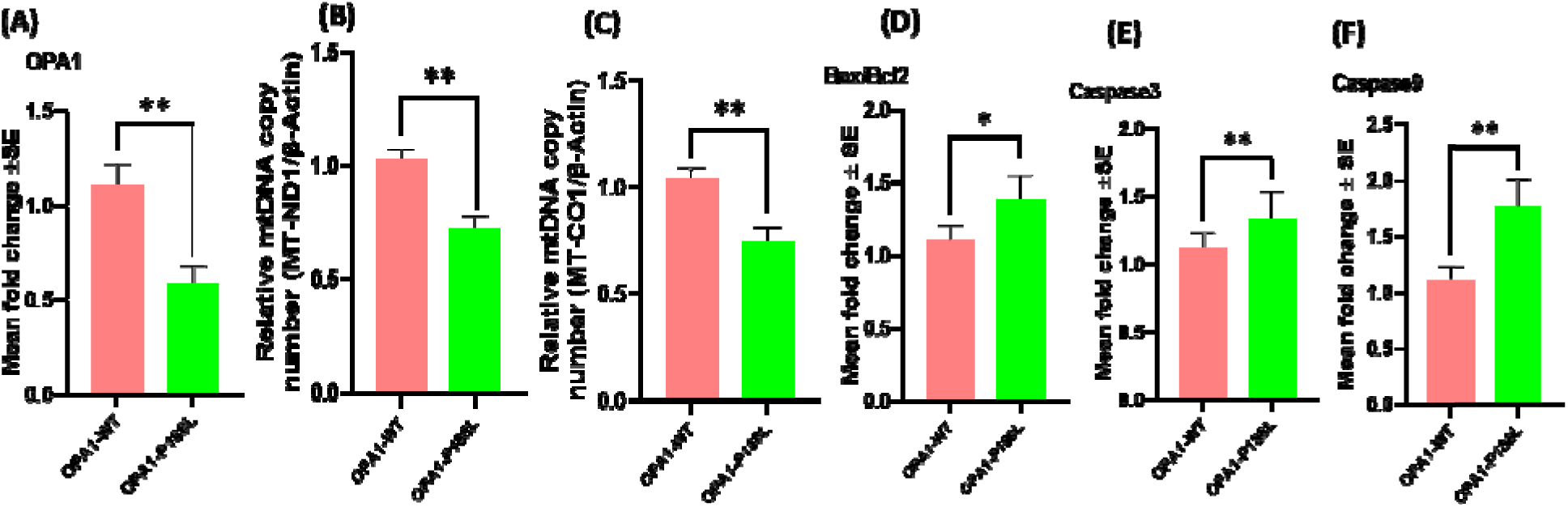
The qPCR detection of the mtDNA copy number and intrinsic apoptotic marker. **A-F** The mRNA levels of ND1, CO1, Bax/Bcl2, Caspase 9 and 3 were determined by qRT-PCR. Three independent experiments for each group. Data are expressed as the mean ± SEM (*P < 0.05, **P<0.01 vs. WT).

### 3.11 Alteration of RNA secondary structure

RNA structure is intimately tied to protein structure and function. Any disruption in RNA folding or sequence can lead to errors in protein synthesis, folding, or regulation. Therefore, we further analysed RNA secondary structure that displayed significant disruptions in RNA folding for OPA1 variant. OPA1 c.563C>T (p.P188L) mutation exhibited pronounced deviations from the WT-RNA secondary structure, suggesting that they might interfere with the normal stability and functional conformation of the RNA molecule. Additionally, a quantitative assessment was performed by calculating the relative entropy [H(wt: mu)] of WT and mutant RNA structures (Table 3). The greater the entropy value provided by remuRNA, the more the structural impact of the variant (Bernhart et al., 2011; Salari et al., 2013). This means, the mutation with higher relative entropy values exhibit greater deviations in RNA structure compared to mutations with lower values. OPA1 mutant variant exhibited entropy value of 2.761, indicating substantial deviation in RNA structure induced by this mutation.

**Table 3.** Analyses of the structural impacts of OPA1 missense variant on RNA.

| OPA1 variant | H(wt:mu) | MFE(mu) | MFE(wt) | dMFE |
| --- | --- | --- | --- | --- |
| P188L | 2.761 | -5.4 | -4.6 | 0.8 |

Abbreviations: H-Relative entropy between wildtype and mutant RNAs, MFE(mu)-minimum free energy (for the mutant sequence), MFE(wt) minimum free energy (for the wildtype sequence); dMFE-difference in MFE between mutant and wildtype

Likewise, the effect of the OPA1 variant on RNA structure is illustrated using multiple graphical approaches, including Circos plots, base pairing probabilities dot plots, differential base pairing probabilities dot plot and RNA accessibility profile analysis (Figure 10). These MutaRNA findings are related to the effect of variant within the RNA snippet affecting the secondary structure and accessibility to its surrounding context.

**Figure 10:**
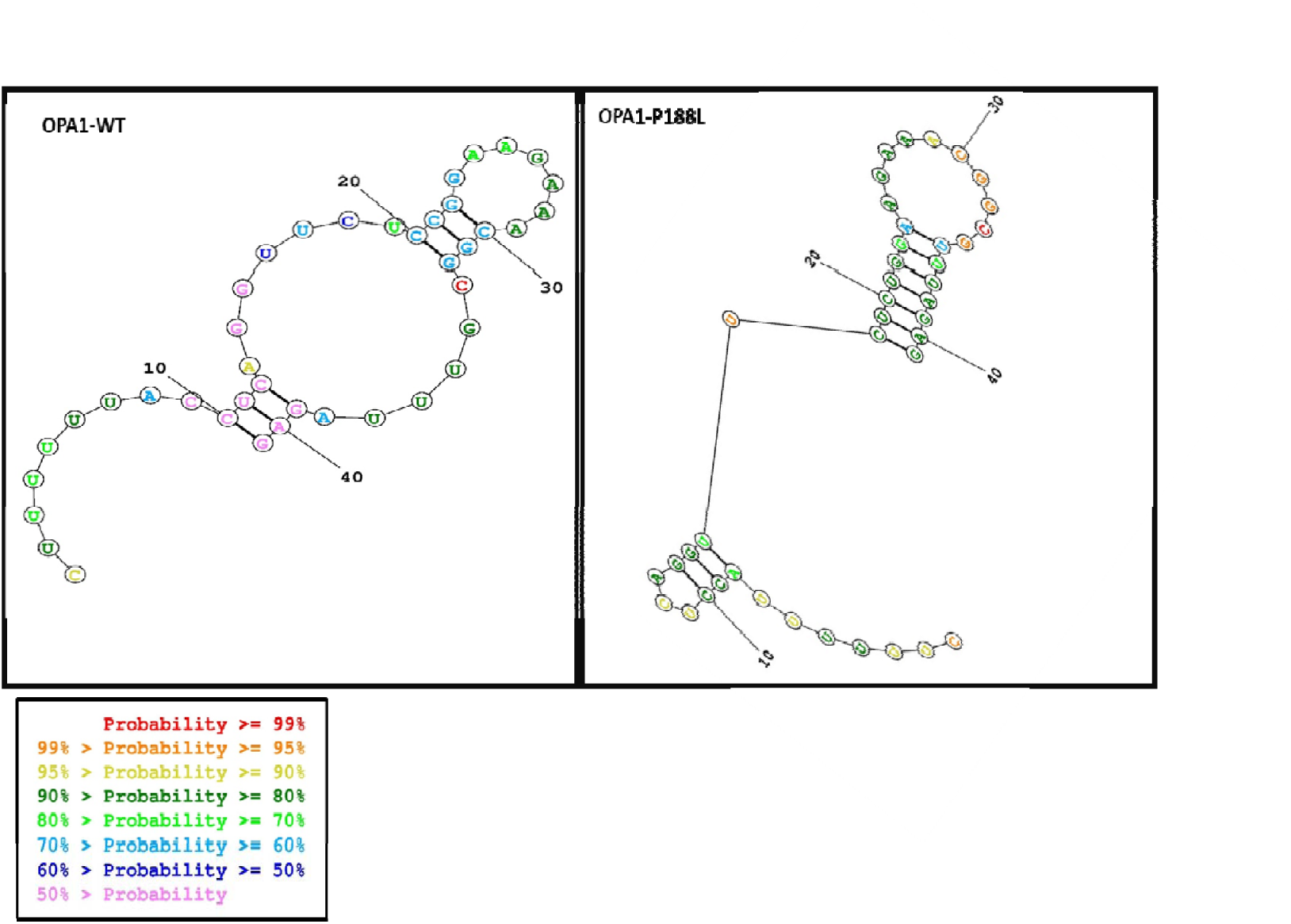
Comparison of Secondary Structures of WT and Mutant (MT) RNA for OPA1 Missense Variant. The WT RNA structure is illustrated on the left, while the mutant RNA structure, altered due to a nucleotide change at the 21st position, is depicted on the right. Base pairing probabilities are represented by varying colours, highlighting structural differences between the two sequences.

We have observed prominent deviation from WT patterns compare to p.P188L indicating significant structural impacts. These base-pair probabilities for WT and mutant (MUT) sequences can also be visualized in heat map dot matrices (figure 11). Consistent alterations in base pairing patterns were observed in variant indicating significant structural changes. This is also evident in the entropy chart and the secondary structures predictions using 2D structure and circos plots base pairing probabilities. The differences in base pairing probabilities between mutant and WT RNA (Δ = Pr(bp in WT) - Pr(bp in MUT)) for OPA1 missense variant are depicted as differential heat map dot matrices (Figure 11) to compare base pairing probabilities between mutant and WT RNA sequences with red indicating increased interaction likelihood and blue indicating weakened base pairs.

**Figure 11:**
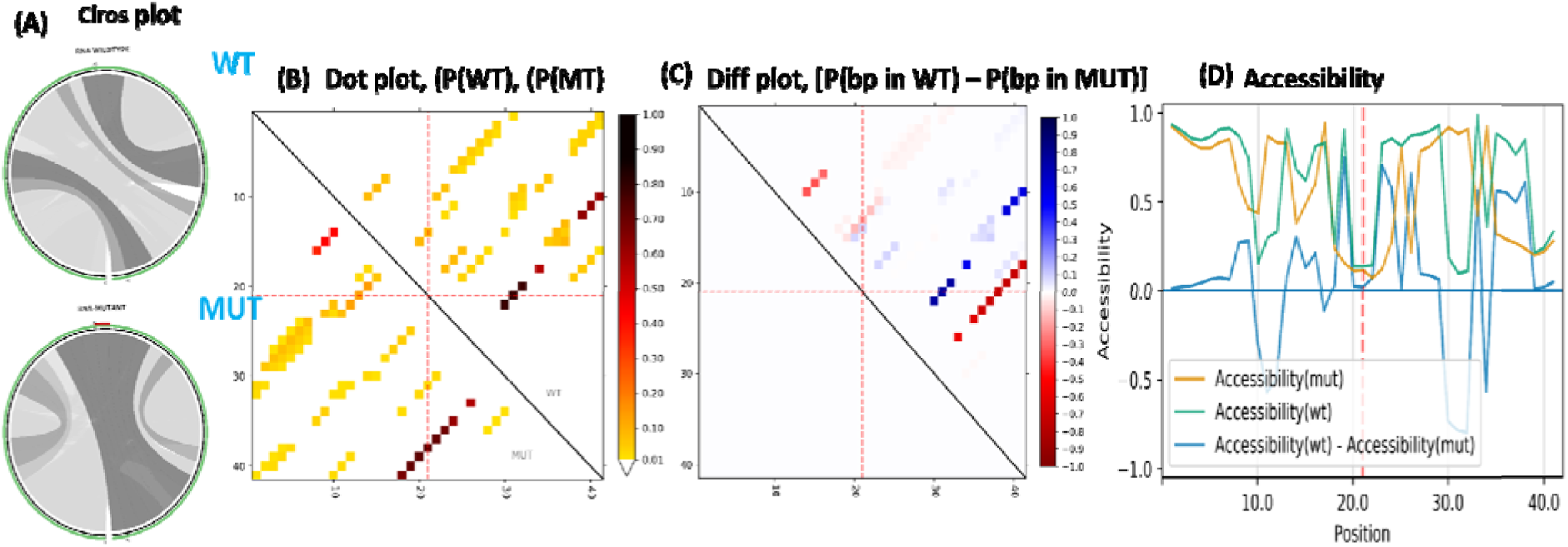
Comparative analysis of RNA structural features of MUT with WT. **(A)**The Circos plots represent the base pairing probability of p(WT) and p(MUT) RNA structures, with WT plot at upper and MUT at lower. The sequence starting from 5’ end at the bottom-left and extending clockwise reaching to the 3’ end. The variant is at position 21 which is highlighted in red at the top of each MUT circos plot. **(B)** The base-pairing of higher probability is illustrated in darker shades of grey. The dot plot is a heatmap-like representation of base-pair probability of p(WT) and p(MUT) structures. The base-pairing of higher probability is indicated in darker dots. **(C)** The differential dot plot depicts the differences in base-pairing probabilities between the WT and MUT RNA [Pr (bp in WT) – Pr (bp in mut)]. The red color dots represent strong base-pairing while the blue color dots highlighting weak base pairing which results due to mutations. **(D)** The column graph demonstrating the accessibility profile in terms of unpaired probabilities of WT and MUT RNA. The blue line indicating the variation in accessibility (WT-MUT) with negative values depicting the positions more prone to be unpaired in the MUT vs WT.

Comparison of accessibility profiles of WT and mutant RNA sequences for OPA1 missense variant (p.P188L) was compared to assess changes in single-stranded-ness and structural dynamics (Figure 11), which is strongly related to its interactions with other proteins or RNAs. The accessibility profile indicates the likelihood of nucleotides being unpaired for each position within the RNA sequences. The blue line illustrates the alteration in accessibility (WT-mut), where negative values signify positions more likely to be unpaired in the mutant compared to the WT. The “negative drops" in the blue differential accessibility profile signify the reduced accessibility in the folded WT compared to its mutant. Which further support RNA secondary structure determination as well as entropy assessment study.

## 4. Discussion

Optic atrophy1 *(OPA1)* gene is a nuclear gene, encoding for a mitochondrial inner membrane-bound protein, localized to chromosome 3q28-29 (HGNC: 8140) comprising 31 exons, that expresses 8 distinct mRNA isoforms. Originally discovered Isoform 1, is translated into a 960 aa long protein with eight specific domains (Figure 2A). Structurally, the OPA1 protein comprises an N-terminal MIS/MTS (mitochondria import/ Targetting sequence) spanning (aa 1-80), followed by a transmembrane domain (aa 96-117), a coiled-coil domain 1 (aa 226-268) that mediates protein–protein interactions, conserved GTPase domain (aa 265-561) drives inner membrane fusion (Belenguer & Pellegrini, 2013), followed by the middle (Stalk) oligomerisation domain (aa 562-800) and Lipid-interacting paddle/LIS: approximately (aa 736–856) support structural stability and lipid-membrane interactions (Praefcke & McMahon, 2004; DeVay et al., 2009) and the C-terminal GTPase effector domain (GED) (aa 800-960) which overlap the coiled-coil 2 domain (aa 940-960) is required to coordinate and enhance GTPase activity (Landes et al., 2010).

The OPA1 protein is a dynamin-related GTPase that plays a central role in mitochondria fusion (Burté et al., 2015). In addition, OPA1 is essential for the assembly and stability of the mitochondrial respiratory chain super-complexes (Olichon et al., 2003), mitochondrial cristae organisation (Frezza et al., 2006) the sequestration of pro-apoptotic cytochrome c molecules within these tight junctions and, importantly, mtDNA maintenance (Elachouri et al., 2011).

In the present study, we have identified a rare missense variation c.532C>T; p.(P188L) in OPA1, detected by WES, which is absent in 100 ethnic-matched heathy control from same geographical location. This variant is present in 5month old male patient exhibiting severe left ventricular dysfunction, having LVEF-21% and enlaged LVIDd = 89 mm, (left ventricular internal diameter in diastole). This variant is absent in different databases such as 1000Genomes, HGMD disease as well as in GenomeAsia 100k and in Indian databases (Table 1). Further, the variation is confined to N-terminal, dynamic central region of OPA1, adjacent to OMA1 clevage site (aa position 195). Multiple bioinformatic tools applying ‘VarCard’ analysis, (including Mutation Taster, SIFT, Polyphen2, LRT-pred, FATHMM, VEST3, Elgen, GenoCanyon etc.) predicted deleterious nature of this variant (Table 2).

A growing body of research has established the pathogenic role of OPA1 mutations, which are primarily associated with Autosomal Dominant Optic Atrophy (ADOA), the most common inherited optic neuropathy(Alexander et al., 2000; Delettre et al., 2000; Marchbank et al., 2002). More than 100 mutations are associated OPA1 gene, causing different types of neuropathy. Approximately 20% of mutation carriers develop a more severe multisystem phenotype, known as DOA-plus syndrome, characterized by chronic progressive external ophthalmoplegia, sensorineural hearing loss, ataxia, myopathy, and peripheral neuropathy(Amati-Bonneau et al., 2008; Yu-Wai-Man et al., 2010). OPA1 mutations have also been associated with Behr-like syndrome, Parkinsonism, and dementia (Marelli et al., 2011; Carelli et al., 2015; Lynch et al., 2017; Del Dotto et al., 2018). Besides these, OPA1 mutations have also been linked to cardiomyopathy together with mitochondrial dysfunction and mtDNA stability (Chen et al., 2012). Notably, the OPA1 p.L534R mutation has been reported to cause lethal infantile encephalopathy, hypertrophic cardiomyopathy, and optic atrophy, accompanied by severe mitochondrial respiratory chain deficiency and marked mitochondrial DNA depletion, further highlighting the critical role of OPA1 in maintaining mitochondrial bioenergetics and tissue homeostasis (Spiegel et al., 2016). Electron microscopy of cardiac tissue from DCM patients exhibited abnormal mitochondrial shape and size as well as loss of cristae structure, which was associated with aberrant OPA1proteolysis (Pawlak et al., 2022). This study was substantiated by a previous report of cardiac specific KO of Yme1l in mice caused OMA1 mediated enhanced OPA1 cleavage triggering mitochondrial fission leading to dilated cardiomyopathy (Wai et al., 2015).

Despite the high expression of OPA1 in the heart, its function in adult cardiomyocytes was first thought to restricted due to the diminished prominence of mitochondrial dynamics in cardiac cells (Dorn et al., 2011; Piquereau et al., 2012). Genetic study has shown that OPA1 is crucial for maintaining cardiac mitochondrial integrity. Homozygous deletion of OPA1 results in embryonic lethality (Davies et al., 2007), whereas Opa1 +/− animals exhibit age-related cardiomyopathy linked to diminished mitochondrial DNA copy number and compromised antioxidant defence (Chen et al., 2012).

The variant identified in this study c.532C>T; p.(P188L) N-terminal region of OPA1 adjacent to the S1 proteolytic cleavage site recognised by OMA1. This region is highly conserved across species, highlighting its functional importance in maintaining OPA1 structure and activity. Variants affecting conserved residues are likely to disrupt the stability, processing, or maturation of OPA1, which may explain the marked reduction observed in both OPA1 protein expression (western blot) and OPA1 transcript levels (∼0.52-fold decrease by qRT-PCR).

Functionally, cells expressing the p.P188L variant exhibited pronounced mitochondrial morphological abnormalities, characterised by a significant increase in intermediate and fragmented mitochondria, indicating impaired mitochondrial inner membrane fusion. Previous studies have demonstrated that OPA1 mutations can produce distinct morphological phenotypes depending on the affected domain. For example, the p.P400A mutation identified in patients with autosomal dominant optic atrophy caused aberrant mitochondrial morphology and network disorganisation (Zhang et al., 2017). Similarly, mutations affecting the GTPase domain (c.870+5G>A and c.889C>T) were associated with elongated mitochondria, whereas mutations in the GED domain (c.2713C>T and c.2818+5G>A) resulted in a predominance of fragmented mitochondria (Cartes-Saavedra et al., 2023).

The phenotype observed in our study was characterized by a predominantly fragmented mitochondrial morphology, suggesting impaired mitochondrial inner membrane fusion. It is plausible that the P188L substitution alters OMA1-mediated proteolytic processing of OPA1, thereby disturbing the balance between the long (L-OPA1) and short (S-OPA1) isoforms. L- OPA1 is required for mitochondrial inner membrane fusion and cristae maintenance, whereas excessive accumulation of S-OPA1 has been associated with mitochondrial fragmentation and fission (Anand et al., 2014; Del Dotto et al., 2018). Notably, recent evidence indicates that GTPase-inactive S-OPA1 can partially localize to mitochondria-associated endoplasmic reticulum membranes (MAMs), suggesting a potential role in ER–mitochondria communication and calcium homeostasis (Anand et al., 2014; Fogo et al., 2024).

OPA1-MUT (P188L)-expressing cells demonstrated a reduction in TMRE fluorescence intensity, indicating significant mitochondrial membrane depolarisation (ΔΨm loss) and impaired mitochondrial function. Consistent with the reduction in ΔΨm, intracellular ATP levels were markedly decreased in P188L-expressing cells compared with OPA1-WT controls, suggesting compromised mitochondrial bioenergetics function. Comprehensive assessment of mitochondrial respiratory function using oxygen consumption rate (OCR) analysis further substantiated these defects. Cells expressing OPA1-P188L exhibited substantial decreases in basal respiration, maximal respiration, ATP-coupled respiration, and spare respiratory capacity. The decreased ATP-linked oxygen consumption rate portion specifically signifies compromised oxidative phosphorylation efficiency, while the decline in spare respiratory capacity implies a restricted capacity of mutant cells to meet heightened energy demands.

These functional abnormalities further supported by *in silico* protein stability predictions via ‘DYNAMUT2, MUPro, DUET, I-Mutant, CUPSTAT, SDM, which predicated that the P188L substitution exerts a destabilizing effect on OPA1. These computational approaches integrate structural, energetic, and evolutionary features to estimate the impact of amino acid substitutions on protein stability. The concordance among these independent prediction tools strengthens the evidence that the p.P188L variant may destabilize the native conformation of OPA1, potentially impairing its role in mitochondrial inner membrane fusion and cristae maintenance, both of which are critical for maintaining cellular energy homeostasis.

Moreover, the superimposed 3D structures of OPA1-P188L mutant versus wild-type show pronounced conformational changes between the two models, as reflected by a root mean square deviation (RMSD) value of 3.5 Å (Figure 3), suggest substantial disruptions in the spatial organisation of the mutant protein, indicating a potential destabilising effect. Such a high RMSD value suggests that the mutation induces remarkable conformational alterations, potentially disrupting the native folding pattern, domain orientation, and interaction interface of OPA1, which may in turn compromise its functional association with partner proteins. Taken together, the diminution of transcript as well as protein level and the functional deficits are likely stem from structural destabilisation of the mutant protein. These data collectively indicate that the P188L mutation significantly undermines mitochondrial bioenergetics integrity, resulting in compromised oxidative phosphorylation, decreased coupling efficiency, and lower cellular energy stores.

In contrast to bioenergetics deficiencies, the present study demonstrated significantly elevated levels of cytosolic Ca² and reactive oxygen species (ROS) in OPA1-MUT (P188L) cells. The mutant cells exhibited increased cytosolic Ca² concentrations together with enhanced ROS generation, indicating disruption of mitochondrial homeostasis. OPA1 is closely associated with the regulation of mitochondrial permeability transition pore (mPTP) susceptibility through its critical role in maintaining cristae junction integrity, mitochondrial inner membrane architecture, and membrane potential. Loss of OPA1 function or excessive OPA1 proteolytic cleavage destabilizes cristae structure, perturbs mitochondrial Ca² handling, and promotes ROS accumulation, thereby increasing susceptibility to Ca² - induced mPTP opening and cytochrome c release, ultimately facilitating apoptotic cell death (Frezza et al., 2006; Papanicolaou et al., 2012; Varanita et al., 2015).

As previously shown in mouse embryonic fibroblasts and HeLa cells, OPA1 depletion disrupted mitochondrial cristae organization and mitochondrial Ca2+ uptake and buffering efficiency, leading to increased oxidative stress and mitochondrial dysfunction (Olichon et al., 2003; Frezza et al., 2006; Cogliati et al., 2013; Cartes-Saavedra et al., 2023). These findings are consistent with our observations and suggest that the OPA1-P188L variant may disrupt mitochondrial Ca2+ homeostasis, resulting in increased cytosolic Ca2+, increased reactive oxygen species (ROS) production, decreased mitochondrial metabolic efficiency and aggravation of oxidative phosphorylation defects.

OPA1 and its yeast homolog Mgm1 are crucial for the organization of the mitochondrial inner membrane and the preservation of cristae, with OPA1 additionally playing a direct role in the maintenance of the mitochondrial genome, comprising mtDNA replication and nucleoid distribution (Griparic et al., 2004; Meeusen et al., 2006; Elachouri et al., 2011). Pathogenic OPA1 mutations have been linked to mtDNA instability, including multiple mtDNA deletions and depletion, in cultured cells and skeletal muscle tissues from subjects with autosomal dominant optic atrophy (ADOA) (Kim et al., 2005; Amati-Bonneau et al., 2008; Liao et al., 2017; Weisschuh et al 2021). In accordance with these reports, the present study showed a reduction of ∼30.6% in mtDNA content in OPA1-mutant cells, indicating defective mitochondrial genome maintenance associated with the OPA1 P188L variant. MtDNA encodes essential subunits of the oxidative phosphorylation (OXPHOS) complexes, and thus mtDNA depletion can affect mitochondrial transcription and translation, disrupt respiratory chain assembly, and thus decrease ATP production (Amati-Bonneau et al., 2008; Chen et al., 2012). These defects may contribute to mitochondrial dysfunction of OPA1- mutant cells together with altered Ca2+ homeostasis and increased oxidative stress.

Importantly, OPA1 is also a key regulator of mitochondrial morphogenesis and apoptosis. OPA1 loss has been shown to cause apoptotic cell death and Bcl-2 overexpression suppresses OPA1 RNAi induced apoptosis, suggesting that mitochondrial fusion is upstream of mitochondrial outer membrane permeabilization (Olichon et al., 2003). Conversely, OPA1 overexpression prevents mitochondrial fission and protects cells from mitochondria-dependent apoptotic death, but not from apoptosis induced by the extrinsic pathway(Frezza et al., 2006). Mechanistically, mitochondrial stress leads to mitochondrial outer membrane permeabilization and cytochrome c release, which binds to Apaf-1 and procaspase-9 to form the apoptosome, resulting in the sequential activation of caspase-9 and caspase-3 and finally nuclear fragmentation (Olichon et al., 2007). Consistent with this mechanism, our study showed increased expression of caspase-9 and caspase-3 in OPA1-P188L mutant cells relative to wild-type controls indicating activation of the intrinsic mitochondrial apoptotic signalling.

OPA1 mRNA has elevated expression during the development and maturation of energy-intensive tissue, such as the heart, brain, skeletal muscle, and retina.(Delettre et al., 2000; Belenguer & Pellegrini, 2013; Tezze et al., 2017). To further investigate the influence of missense mutations in OPA1 on RNA secondary structure, various computational approaches have been utilised. The prediction of RNA secondary structure indicates that RNA folding and stability are dramatically affected by the mutation, suggesting potential functional consequences at the molecular level. The structural differences between wild-type and mutant RNA are assessed using relative entropy calculations to evaluate the degree of mutation-induced alterations. To corroborate these findings, various visualisation techniques are utilised, including Circos plots, base pair probability dot plots, and differential base-pairing probability dot plots (Figure 10). Furthermore, accessibility profiles, denoting the likelihood of particular nucleotides remaining unpaired, were examined for both wild-type and mutant RNA sequences. The variations in accessibility profiles provided insights into how mutations may modify RNA interactions with proteins or other RNAs. OPA1 variants that impair splice-site RNA structure are viable targets for precision medicine, as RNA-based therapeutic approaches can correct abnormal splicing, restore functional transcript equilibrium, maintain mitochondrial integrity, and reduce apoptotic vulnerability without causing permanent genomic changes (Jüschke et al., 2021). This investigation highlights the influence of variation on RNA structure and function, offering insights into the molecular pathways associated with disease aetiology (Salari et al., 2013).

## Conclusion

In a nutshell, our results strongly suggest that OPA1 is required for maintaining mitochondrial integrity, bioenergetic homeostasis, and cell survival in cardiomyocytes. The combined *in-vitro* and i*n-silico* studies suggest that the OPA1 (P188L) variation results in decreased stability of OPA1 protein and structural conformational changes that affect its functional interactions and proteolytic processing, thus impairing its function in mitochondrial inner membrane fusion and cristae maintenance. These perturbations are linked with disrupted mitochondrial dynamics, aberrant calcium homeostasis, defective mitochondrial genome maintenance, compromised oxidative phosphorylation, increased oxidative stress and activation of intrinsic apoptotic signalling. Such coordinated defects are expected to increase susceptibility of cardiomyocytes to mitochondrial injury and pathological cardiac remodelling. Our study supports the contribution of OPA1-mediated mitochondrial dysfunction to the pathogenesis of dilated cardiomyopathy, and highlights OPA1 as a critical regulator of cardiac mitochondrial function given its recognized crucial role in mitochondrial quality control and energy homeostasis. These results provide a foundation for future mechanistic studies and the development of therapeutic stratecgies to target OPA1 stability, mitochondrial dynamics and mitochondrial quality-control pathways in cardiomyopathy.

## Acknowledgment

We sincerely thank the patients and their families for their invaluable participation in this study. We also acknowledge Prof. Daniel Linseman (University of Denver, Denver, CO) for providing the full-length OPA1 clone. We thank the Sophisticated Analytical Instrument Facility (SAIF), AIIMS, New Delhi, for TEM facilities, and the Sophisticated Analytical & Technical Help Institute (SATHI), Banaras Hindu University, Varanasi, for access to the laser-scanning super-resolution microscopy system. We are grateful to Prof. D. Das, Department of Biochemistry, Institute of Medical Sciences, Banaras Hindu University, Varanasi, for providing access to the Oxygraph-2k system (Oroboros Instruments, Innsbruck, Austria). This work was supported by the Department of Biotechnology (DBT), Government of India (BT/PR12369/MED/12/678/2014), and by an ICMR Senior Research Fellowship awarded to Ms.Mohini Gupta (F. No. 2021-11036/Proteomics-BMS).

## Author contributions

Bhagyalaxmi Mohapatra: Conceptualization, Supervision, Investigation, Validation, Resources, Writing-Review & editing, Mohini Gupta: Data curation, Validation, Software, Methodology, Formal analysis, Visualization, Writing- Original draft preparation. Amrita Mukhopadhyay for methology. Ashok Kumar: for identifying and enrolling patients for the study.

## References

1. Alavi, M. V. (2021). OMA1—An integral membrane protease? Biochimica et Biophysica Acta (BBA) - Proteins and Proteomics, 1869(2), 140558. 10.1016/j.bbapap.2020.140558

2. Alexander, C., Votruba, M., Pesch, U. E. A., Thiselton, D. L., Mayer, S., Rodriguez, M., Kellner, U., Leo-Kottler, B., Auburger, G., Bhattacharya, S. S., & Wissinger, B. (2000). OPA1, encoding a dynamin-related GTPase, is mutated in autosomal dominant optic atrophy linked to chromosome 3q28.

3. Amati-Bonneau, P., Valentino, M. L., Reynier, P., Gallardo, M. E., Bornstein, B., Boissiere, A., Campos, Y., Rivera, H., De La Aleja, J. G., Carroccia, R., Iommarini, L., Labauge, P., Figarella-Branger, D., Marcorelles, P., Furby, A., Beauvais, K., Letournel, F., Liguori, R., La Morgia, C., … Carelli, V. (2008). OPA1 mutations induce mitochondrial DNA instability and optic atrophy “plus” phenotypes. Brain, 131(2), 338–351. 10.1093/brain/awm298

4. Anand, R., Wai, T., Baker, M. J., Kladt, N., Schauss, A. C., Rugarli, E., & Langer, T. (2014). The *i* -AAA protease YME1L and OMA1 cleave OPA1 to balance mitochondrial fusion and fission. Journal of Cell Biology, 204(6), 919–929. 10.1083/jcb.201308006

5. Baker, M. J., Lampe, P. A., Stojanovski, D., Korwitz, A., Anand, R., Tatsuta, T., & Langer, T. (2014). Stress-induced OMA1 activation and autocatalytic turnover regulate OPA1-dependent mitochondrial dynamics. The EMBO Journal, 33(6), 578–593. 10.1002/embj.201386474

6. Ban, T., Ishihara, T., Kohno, H., Saita, S., Ichimura, A., Maenaka, K., Oka, T., Mihara, K., & Ishihara, N. (2017). Molecular basis of selective mitochondrial fusion by heterotypic action between OPA1 and cardiolipin. Nature Cell Biology, 19(7), 856–863. 10.1038/ncb3560

7. Barrera, M., Koob, S., Dikov, D., Vogel, F., & Reichert, A. S. (2016). OPA1 functionally interacts with MIC60 but is dispensable for crista junction formation. FEBS Letters, 590(19), 3309–3322. 10.1002/1873-3468.12384

8. Belenguer, P., & Pellegrini, L. (2013a). The dynamin GTPase OPA1: More than mitochondria? Biochimica et Biophysica Acta (BBA) - Molecular Cell Research, 1833(1), 176–183. 10.1016/j.bbamcr.2012.08.004

9. Belenguer, P., & Pellegrini, L. (2013b). The dynamin GTPase OPA1: More than mitochondria? Biochimica et Biophysica Acta (BBA) - Molecular Cell Research, 1833(1), 176–183. 10.1016/j.bbamcr.2012.08.004

10. Bellaousov, S., Reuter, J. S., Seetin, M. G., & Mathews, D. H. (2013). RNAstructure: Web servers for RNA secondary structure prediction and analysis. Nucleic Acids Research, 41(W1), W471–W474. 10.1093/nar/gkt290

11. Bernhart, S. H., Mückstein, U., & Hofacker, I. L. (2011). RNA Accessibility in cubic time. Algorithms for Molecular Biology, 6(1), 3. 10.1186/1748-7188-6-3

12. Burté, F., Carelli, V., Chinnery, P. F., & Yu-Wai-Man, P. (2015). Disturbed mitochondrial dynamics and neurodegenerative disorders. Nature Reviews Neurology, 11(1), 11–24. 10.1038/nrneurol.2014.228

13. Carelli, V., Musumeci, O., Caporali, L., Zanna, C., La Morgia, C., Del Dotto, V., Porcelli, A. M., Rugolo, M., Valentino, M. L., Iommarini, L., Maresca, A., Barboni, P., Carbonelli, M., Trombetta, C., Valente, E. M., Patergnani, S., Giorgi, C., Pinton, P., Rizzo, G., … Zeviani, M. (2015). Syndromic parkinsonism and dementia associated with *OPA 1* missense mutations. Annals of Neurology, 78(1), 21–38. 10.1002/ana.24410

14. Cartes-Saavedra, B., Lagos, D., Macuada, J., Arancibia, D., Burté, F., Sjöberg-Herrera, M. K., Andrés, M. E., Horvath, R., Yu-Wai-Man, P., Hajnóczky, G., & Eisner, V. (2023). *OPA1* disease-causing mutants have domain-specific effects on mitochondrial ultrastructure and fusion. Proceedings of the National Academy of Sciences, 120(12), e2207471120. 10.1073/pnas.2207471120

15. Chen, H., Chomyn, A., & Chan, D. C. (2005). Disruption of Fusion Results in Mitochondrial Heterogeneity and Dysfunction. Journal of Biological Chemistry, 280(28), 26185– 26192. 10.1074/jbc.M503062200

16. Chen, L., Liu, T., Tran, A., Lu, X., Tomilov, A. A., Davies, V., Cortopassi, G., Chiamvimonvat, N., Bers, D. M., Votruba, M., & Knowlton, A. A. (2012). OPA1 Mutation and Late Onset Cardiomyopathy: Mitochondrial Dysfunction and mtDNA Instability. Journal of the American Heart Association, 1(5), e003012. 10.1161/JAHA.112.003012

17. Cogliati, S., Frezza, C., Soriano, M. E., Varanita, T., Quintana-Cabrera, R., Corrado, M., Cipolat, S., Costa, V., Casarin, A., Gomes, L. C., Perales-Clemente, E., Salviati, L., Fernandez-Silva, P., Enriquez, J. A., & Scorrano, L. (2013). Mitochondrial Cristae Shape Determines Respiratory Chain Supercomplexes Assembly and Respiratory Efficiency. Cell, 155(1), 160–171. 10.1016/j.cell.2013.08.032

18. Davies, V. J., Hollins, A. J., Piechota, M. J., Yip, W., Davies, J. R., White, K. E., Nicols, P. P., Boulton, M. E., & Votruba, M. (2007). Opa1 deficiency in a mouse model of autosomal dominant optic atrophy impairs mitochondrial morphology, optic nerve structure and visual function. Human Molecular Genetics, 16(11), 1307–1318. 10.1093/hmg/ddm079

19. Del Dotto, V., Fogazza, M., Carelli, V., Rugolo, M., & Zanna, C. (2018). Eight human OPA1 isoforms, long and short: What are they for? Biochimica et Biophysica Acta (BBA) - Bioenergetics, 1859(4), 263–269. 10.1016/j.bbabio.2018.01.005

20. Del Dotto, V., Fogazza, M., Musiani, F., Maresca, A., Aleo, S. J., Caporali, L., La Morgia, C., Nolli, C., Lodi, T., Goffrini, P., Chan, D., Carelli, V., Rugolo, M., Baruffini, E., & Zanna, C. (2018). Deciphering OPA1 mutations pathogenicity by combined analysis of human, mouse and yeast cell models. Biochimica et Biophysica Acta (BBA) - Molecular Basis of Disease, 1864(10), 3496–3514. 10.1016/j.bbadis.2018.08.004

21. Delettre, C., Lenaers, G., Griffoin, J.-M., Gigarel, N., Lorenzo, C., Belenguer, P., Pelloquin, L., Grosgeorge, J., Turc-Carel, C., Perret, E., Astarie-Dequeker, C., Lasquellec, L., Arnaud, B., Ducommun, B., Kaplan, J., & Hamel, C. P. (2000). Nuclear gene OPA1, encoding a mitochondrial dynamin- related protein, is mutated in dominant optic atrophy.

22. DeVay, R. M., Dominguez-Ramirez, L., Lackner, L. L., Hoppins, S., Stahlberg, H., & Nunnari, J. (2009). Coassembly of Mgm1 isoforms requires cardiolipin and mediates mitochondrial inner membrane fusion. Journal of Cell Biology, 186(6), 793–803. 10.1083/jcb.200906098

23. Dorn, G. W., Clark, C. F., Eschenbacher, W. H., Kang, M.-Y., Engelhard, J. T., Warner, S. J., Matkovich, S. J., & Jowdy, C. C. (2011). MARF and Opa1 Control Mitochondrial and Cardiac Function in Drosophila. Circulation Research, 108(1), 12–17. 10.1161/CIRCRESAHA.110.236745

24. Elachouri, G., Vidoni, S., Zanna, C., Pattyn, A., Boukhaddaoui, H., Gaget, K., Yu-Wai-Man, P., Gasparre, G., Sarzi, E., Delettre, C., Olichon, A., Loiseau, D., Reynier, P., Chinnery, P. F., Rotig, A., Carelli, V., Hamel, C. P., Rugolo, M., & Lenaers, G. (2011). OPA1 links human mitochondrial genome maintenance to mtDNA replication and distribution. Genome Research, 21(1), 12–20. 10.1101/gr.108696.110

25. Fogo, G. M., Raghunayakula, S., Emaus, K. J., Torres Torres, F. J., Wider, J. M., & Sanderson, T. H. (2024). Mitochondrial membrane potential and oxidative stress interact to regulate Oma1 dependent processing of Opa1 and mitochondrial dynamics. The FASEB Journal, 38(18), e70066. 10.1096/fj.202400313R

26. Frezza, C., Cipolat, S., Martins De Brito, O., Micaroni, M., Beznoussenko, G. V., Rudka, T., Bartoli, D., Polishuck, R. S., Danial, N. N., De Strooper, B., & Scorrano, L. (2006). OPA1 Controls Apoptotic Cristae Remodeling Independently from Mitochondrial Fusion. Cell, 126(1), 177–189. 10.1016/j.cell.2006.06.025

27. Garcia, I., Calderon, F., La Torre, P. D., Vallier, S. St., Rodriguez, C., Agarwala, D., Keniry, M., Innis-Whitehouse, W., & Gilkerson, R. (2021). Mitochondrial OPA1 cleavage is reversibly activated by differentiation of H9c2 cardiomyoblasts. Mitochondrion, 57, 88–96. 10.1016/j.mito.2020.12.007

28. Griparic, L., Van Der Wel, N. N., Orozco, I. J., Peters, P. J., & Van Der Bliek, A. M. (2004). Loss of the Intermembrane Space Protein Mgm1/OPA1 Induces Swelling and Localized Constrictions along the Lengths of Mitochondria. Journal of Biological Chemistry, 279(18), 18792–18798. 10.1074/jbc.M400920200

29. Jiang, P., Wang, M., Xue, L., Xiao, Y., Yu, J., Wang, H., Yao, J., Liu, H., Peng, Y., Liu, H., Li, H., Chen, Y., & Guan, M.-X. (2016). A Hypertension-Associated tRNA^Ala^ Mutation Alters tRNA Metabolism and Mitochondrial Function. Molecular and Cellular Biology, 36(14), 1920–1930. 10.1128/MCB.00199-16

30. Jüschke, C., Klopstock, T., Catarino, C. B., Owczarek-Lipska, M., Wissinger, B., & Neidhardt, J. (2021). Autosomal dominant optic atrophy: A novel treatment for OPA1 splice defects using U1 snRNA adaption. Molecular Therapy - Nucleic Acids, 26, 1186–1197. 10.1016/j.omtn.2021.10.019

31. Kim, J. Y., Hwang, J.-M., Ko, H. S., Seong, M.-W., Park, B.-J., & Park, S. S. (2005). Mitochondrial DNA content is decreased in autosomal dominant optic atrophy. Neurology, 64(6), 966–972. 10.1212/01.WNL.0000157282.76715.B1

32. Kushnareva, Y. E., Gerencser, A. A., Bossy, B., Ju, W.-K., White, A. D., Waggoner, J., Ellisman, M. H., Perkins, G., & Bossy-Wetzel, E. (2013). Loss of OPA1 disturbs cellular calcium homeostasis and sensitizes for excitotoxicity. Cell Death & Differentiation, 20(2), 353–365. 10.1038/cdd.2012.128

33. Landes, T., Leroy, I., Bertholet, A., Diot, A., Khosrobakhsh, F., Daloyau, M., Davezac, N., Miquel, M.-C., Courilleau, D., & Guillou, E. (2010). OPA1 (dys)functions. Seminars in Cell & Developmental Biology, 21(6), 593–598. 10.1016/j.semcdb.2009.12.012

34. Lee, H., Smith, S. B., & Yoon, Y. (2017). The short variant of the mitochondrial dynamin OPA1 maintains mitochondrial energetics and cristae structure. Journal of Biological Chemistry, 292(17), 7115–7130. 10.1074/jbc.M116.762567

35. Liao, C., Ashley, N., Diot, A., Morten, K., Phadwal, K., Williams, A., Fearnley, I., Rosser, L., Lowndes, J., Fratter, C., Ferguson, D. J. P., Vay, L., Quaghebeur, G., Moroni, I., Bianchi, S., Lamperti, C., Downes, S. M., Sitarz, K. S., Flannery, P. J., … Poulton, J. (2017). Dysregulated mitophagy and mitochondrial organization in optic atrophy due to *OPA1* mutations. Neurology, 88(2), 131–142. 10.1212/WNL.0000000000003491

36. Lynch, D. S., Loh, S. H. Y., Harley, J., Noyce, A. J., Martins, L. M., Wood, N. W., Houlden, H., & Plun-Favreau, H. (2017). Nonsyndromic Parkinson disease in a family with autosomal dominant optic atrophy due to *OPA1* mutations. Neurology Genetics, 3(5), e188. 10.1212/NXG.0000000000000188

37. MacVicar, T., & Langer, T. (2016). OPA1 processing in cell death and disease – the long and short of it. Journal of Cell Science, 129(12), 2297–2306. 10.1242/jcs.159186

38. Maftah, A., Petit, J. M., Ratinaud, M.-H., & Julien, R. (1989). 10-N Nonyl-Acridine Orange: A fluorescent probe which stains mitochondria independently of their energetic state. Biochemical and Biophysical Research Communications, 164(1), 185–190. 10.1016/0006-291X(89)91700-2

39. Marchbank, N. J., Craig, J. E., Leek, J. P., Toohey, M., Churchill, A. J., Markham, A. F., Mackey, D. A., Toomes, C., & Inglehearn, C. F. (2002). Deletion of the *OPA1* gene in a dominant optic atrophy family: Evidence that haploinsufficiency is the cause of disease. Journal of Medical Genetics, 39(8), e47–e47. 10.1136/jmg.39.8.e47

40. Marelli, C., Amati-Bonneau, P., Reynier, P., Layet, V., Layet, A., Stevanin, G., Brissaud, E., Bonneau, D., Durr, A., & Brice, A. (2011). Heterozygous OPA1 mutations in Behr syndrome. Brain, 134(4), e169–e169. 10.1093/brain/awq306

41. Meeusen, S., DeVay, R., Block, J., Cassidy-Stone, A., Wayson, S., McCaffery, J. M., & Nunnari, J. (2006). Mitochondrial Inner-Membrane Fusion and Crista Maintenance Requires the Dynamin-Related GTPase Mgm1. Cell, 127(2), 383–395. 10.1016/j.cell.2006.09.021

42. Miladi, M., Raden, M., Diederichs, S., & Backofen, R. (2020). MutaRNA: Analysis and visualization of mutation-induced changes in RNA structure. Nucleic Acids Research, 48(W1), W287–W291. 10.1093/nar/gkaa331

43. Noone, J., O’Gorman, D. J., & Kenny, H. C. (2022). OPA1 regulation of mitochondrial dynamics in skeletal and cardiac muscle. Trends in Endocrinology & Metabolism, 33(10), 710–721. 10.1016/j.tem.2022.07.003

44. Olichon, A., Baricault, L., Gas, N., Guillou, E., Valette, A., Belenguer, P., & Lenaers, G. (2003). Loss of OPA1 Perturbates the Mitochondrial Inner Membrane Structure and Integrity, Leading to Cytochrome c Release and Apoptosis. Journal of Biological Chemistry, 278(10), 7743–7746. 10.1074/jbc.C200677200

45. Olichon, A., ElAchouri, G., Baricault, L., Delettre, C., Belenguer, P., & Lenaers, G. (2007). OPA1 alternate splicing uncouples an evolutionary conserved function in mitochondrial fusion from a vertebrate restricted function in apoptosis. Cell Death & Differentiation, 14(4), 682–692. 10.1038/sj.cdd.4402048

46. Papanicolaou, K. N., Kikuchi, R., Ngoh, G. A., Coughlan, K. A., Dominguez, I., Stanley, W. C., & Walsh, K. (2012). Mitofusins 1 and 2 Are Essential for Postnatal Metabolic Remodeling in Heart. Circulation Research, 111(8), 1012–1026. 10.1161/CIRCRESAHA.112.274142

47. Pawlak, A., Gewartowska, M., Przybylski, M., Kuffner, M., Wiligórska, D., Gil, R., Król, Z., & Frontczak-Baniewicz, M. (2022). Ultrastructural Changes in Mitochondria in Patients with Dilated Cardiomyopathy and Parvovirus B19 Detected in Heart Tissue without Myocarditis. Journal of Personalized Medicine, 12(2), 177. 10.3390/jpm12020177

48. Pernas, L., & Scorrano, L. (2016). Mito-Morphosis: Mitochondrial Fusion, Fission, and Cristae Remodeling as Key Mediators of Cellular Function. Annual Review of Physiology, 78(1), 505–531. 10.1146/annurev-physiol-021115-105011

49. Petit, J., Maftah, A., Ratinaud, M., & Julien, R. (1992). 10 *N* Nonyl acridine orange interacts with cardiolipin and allows the quantification of this phospholipid in isolated mitochondria. European Journal of Biochemistry, 209(1), 267–273. 10.1111/j.1432-1033.1992.tb17285.x

50. Piquereau, J., Caffin, F., Novotova, M., Lemaire, C., Veksler, V., Garnier, A., Ventura-Clapier, R., & Joubert, F. (2013). Mitochondrial dynamics in the adult cardiomyocytes: Which roles for a highly specialized cell? Frontiers in Physiology, 4. 10.3389/fphys.2013.00102

51. Piquereau, J., Caffin, F., Novotova, M., Prola, A., Garnier, A., Mateo, P., Fortin, D., Huynh, L. H., Nicolas, V., Alavi, M. V., Brenner, C., Ventura-Clapier, R., Veksler, V., & Joubert, F. (2012). Down-regulation of OPA1 alters mouse mitochondrial morphology, PTP function, and cardiac adaptation to pressure overload. Cardiovascular Research, 94(3), 408–417. 10.1093/cvr/cvs117

52. Praefcke, G. J. K., & McMahon, H. T. (2004). The dynamin superfamily: Universal membrane tubulation and fission molecules? Nature Reviews Molecular Cell Biology, 5(2), 133–147. 10.1038/nrm1313

53. Riss, T. L., Moravec, R. A., Niles, A. L., Benink, H. A., Worzella, T. J., & Minor, L. (2013). Cell Viability Assays.

54. Salari, R., Kimchi-Sarfaty, C., Gottesman, M. M., & Przytycka, T. M. (2013). Sensitive measurement of single-nucleotide polymorphism-induced changes of RNA conformation: Application to disease studies. Nucleic Acids Research, 41(1), 44–53. 10.1093/nar/gks1009

55. Shahrestani, P., Leung, H.-T., Le, P. K., Pak, W. L., Tse, S., Ocorr, K., & Huang, T. (2009). Heterozygous Mutation of Drosophila Opa1 Causes the Development of Multiple Organ Abnormalities in an Age-Dependent and Organ-Specific Manner. PLoS ONE, 4(8), e6867. 10.1371/journal.pone.0006867

56. Spiegel, R., Saada, A., Flannery, P. J., Burté, F., Soiferman, D., Khayat, M., Eisner, V., Vladovski, E., Taylor, R. W., Bindoff, L. A., Shaag, A., Mandel, H., Schuler-Furman, O., Shalev, S. A., Elpeleg, O., & Yu-Wai-Man, P. (2016). Fatal infantile mitochondrial encephalomyopathy, hypertrophic cardiomyopathy and optic atrophy associated with a homozygous *OPA1* mutation. Journal of Medical Genetics, 53(2), 127–131. 10.1136/jmedgenet-2015-103361

57. Tezze, C., Romanello, V., Desbats, M. A., Fadini, G. P., Albiero, M., Favaro, G., Ciciliot, S., Soriano, M. E., Morbidoni, V., Cerqua, C., Loefler, S., Kern, H., Franceschi, C., Salvioli, S., Conte, M., Blaauw, B., Zampieri, S., Salviati, L., Scorrano, L., & Sandri, M. (2017). Age-Associated Loss of OPA1 in Muscle Impacts Muscle Mass, Metabolic Homeostasis, Systemic Inflammation, and Epithelial Senescence. Cell Metabolism, 25(6), 1374–1389.e6. 10.1016/j.cmet.2017.04.021

58. Tobacyk, J., Parajuli, N., Shrum, S., Crow, J. P., & MacMillan-Crow, L. A. (2019). The first direct activity assay for the mitochondrial protease OMA1. Mitochondrion, 46, 1–5. 10.1016/j.mito.2019.03.001

59. Varanita, T., Soriano, M. E., Romanello, V., Zaglia, T., Quintana-Cabrera, R., Semenzato, M., Menabò, R., Costa, V., Civiletto, G., Pesce, P., Viscomi, C., Zeviani, M., Di Lisa, F., Mongillo, M., Sandri, M., & Scorrano, L. (2015). The Opa1-Dependent Mitochondrial Cristae Remodeling Pathway Controls Atrophic, Apoptotic, and Ischemic Tissue Damage. Cell Metabolism, 21(6), 834–844. 10.1016/j.cmet.2015.05.007

60. Votruba, M. (1998). Clinical Features in Affected Individuals From 21 Pedigrees With Dominant Optic Atrophy. Archives of Ophthalmology, 116(3), 351. 10.1001/archopht.116.3.351

61. Wai, T., García-Prieto, J., Baker, M. J., Merkwirth, C., Benit, P., Rustin, P., Rupérez, F. J., Barbas, C., Ibañez, B., & Langer, T. (2015). Imbalanced OPA1 processing and mitochondrial fragmentation cause heart failure in mice. Science, 350(6265), aad0116. 10.1126/science.aad0116

62. Wang, X. (2001). The expanding role of mitochondria in apoptosis. GENES & DEVELOPMENT 15:2922–2933, by Cold Spring Harbor Laboratory Press ISSN 0890-9369/01

63. Winyard, P. G., Ryan, B., Eggleton, P., Nissim, A., Taylor, E., Lo Faro, M. L., Burkholz, T., Szabó-Taylor, K. E., Fox, B., Viner, N., Haigh, R. C., Benjamin, N., Jones, A. M., & Whiteman, M. (2011). Measurement and meaning of markers of reactive species of oxygen, nitrogen and sulfur in healthy human subjects and patients with inflammatory joint disease. Biochemical Society Transactions, 39(5), 1226–1232. 10.1042/BST0391226

64. Yu-Wai-Man, P., Griffiths, P. G., Gorman, G. S., Lourenco, C. M., Wright, A. F., Auer-Grumbach, M., Toscano, A., Musumeci, O., Valentino, M. L., Caporali, L., Lamperti, C., Tallaksen, C. M., Duffey, P., Miller, J., Whittaker, R. G., Baker, M. R., Jackson, M. J., Clarke, M. P., Dhillon, B., … Chinnery, P. F. (2010). Multi-system neurological disease is common in patients with OPA1 mutations. Brain, 133(3), 771– 786. 10.1093/brain/awq007

65. Zerem, A., Yosovich, K., Rappaport, Y. C., Libzon, S., Blumkin, L., Ben-Sira, L., Lev, D., & Lerman-Sagie, T. (2019). Metabolic stroke in a patient with bi-allelic OPA1 mutations. Metabolic Brain Disease, 34(4), 1043–1048. 10.1007/s11011-019-00415-2

66. Zhang, J., Liu, X., Liang, X., Lu, Y., Zhu, L., Fu, R., Ji, Y., Fan, W., Chen, J., Lin, B., Yuan, Y., Jiang, P., Zhou, X., & Guan, M.-X. (2017). A novel ADOA-associated OPA1 mutation alters the mitochondrial function, membrane potential, ROS production and apoptosis. Scientific Reports, 7(1), 5704. 10.1038/s41598-017-05571-y

67. Zhang, K., Li, H., & Song, Z. (2014). Membrane depolarization activates the mitochondrial protease OMA 1 by stimulating self cleavage. EMBO Reports, 15(5), 576–585. 10.1002/embr.201338240

